# Human ERG Oncoprotein Represses *Chip/LDB1* LIM-Domain Binding Gene in *Drosophila*

**DOI:** 10.1101/2022.06.05.494862

**Authors:** Mahima Bharti, Anjali Bajpai, Umanshi Rautela, Nishat Manzar, Bushra Ateeq, Pradip Sinha

## Abstract

ERG oncoprotein, a master transcription factor, targets diverse arrays of genes in different cancers. Identifying oncogenically relevant ones from these ERG targets, however, is challenging. Here we show that heterologous ERG disrupts a LIM-homeodomain (LIM-HD) complex, Chip-Tailup, in *Drosophila*. In the posterior thorax (notum) primordium, ERG-induced upregulation of *E(z)/EZH2* trimethylates histones in *Chip* promoter. A consequent loss of the Chip-Tailup complex releases repression of N-Wg signaling in the notum, inducing *de novo* wings and, alternatively, carcinogenesis of ERG-expressing notal cells displaying loss of Lgl tumor suppressor. ERG-induced developmental or oncogenic fallouts are abrogated upon gain of Chip, N, or E(z) loss, besides Wg ligand sequestration. ERG-positive prostate cancer (PCa) cells, too, display suppression of mammalian homolog of *Drosophila Chip*, *LIM Domain Binding1, LDB1*. Deep homology in gene regulatory networks, like that of Chip-Tup complex, thus help prioritize identification of functionally relevant targets of human oncoproteins in *Drosophila*.

**Highlights:**

- Human ERG suppresses *Chip*, a LIM-domain binding, LDB gene in *Drosophila* via E(z)
- ERG-mediated Chip loss induces ectopic Wg morphogen signaling in the notum primordium
- Chip gain suppresses ERG-induced Wg morphogen and tumor progression in *lgl* clones
- ERG-positive human PCa cell lines show downregulation of a *Chip* homolog, *LDB1*

**In brief:** Mammalian ERG oncoprotein displays a diverse and perplexing range of targets in different cancers. By driving ERG in *Drosophila* developing appendages, Bharti et al. reveal its repression of a LIM-domain coding gene, *Chip/LDB1*. ERG-positive prostate cancer cells, too, display *Chip/LDB1* repression. Deep homology across phylogeny thus helps uncover oncoprotein targets.

## Introduction

The conservation of genes regulating pattern formation in animals across distant phylogenies reveals deep homology (Shubin et al., 2009). Shared genetic tool kits ranging from cellular signaling pathways, homeotic selectors, or master transcription factors underpin such deep homologies (Carroll et al., 2013). Apart from these well-known genetic tool kits, LIM-homeodomain (LIM-HD) transcription factors are also conserved across the animal kingdom (Larroux et al., 2008). Examples of this class of LIM-HD transcription factors include Tailup (Tup) and Apterous (Ap) in *Drosophila;* their mammalian homologs being Islet (Isl) and LIM-homeobox (Lhx), respectively [for reviews, see (Bach, 2000; Jurata and Gill, 2001)]. LIM-HD transcription factors are activated by forming tetrameric complexes with transcription co-factor, LIM-domain binding (LDB) proteins such as Chip, in *Drosophila* (Bach, 2000) and its homolog, LIM Domain Binding1, LDB1, in mammals [reviewed in (Bronstein and Segal, 2011)]. Conservation of LIM-HD complexes across phylogeny (Larroux et al., 2008) also underscores the pervasive nature of deep homology in pattern formation from insects (Jurata and Gill, 2001) to mammals (Kawakami et al., 2011; Narkis et al., 2012).

During mammalian development, expression of ERG (<u>E</u>TS-<u>R</u>elated <u>G</u>ene) is seen in endothelial cells and primordia of organs of mesodermal lineage: for instance, developing kidney, urogenital tract, hematopoietic cells, cartilage, and neural crest cells (Birdsey et al., 2008; Iwamoto et al., 2000). In the adults, ERG expression continues in the cells of endothelial lineage (Mohamed et al., 2010; Vlaeminck-Guillem et al., 2000), while its expression is not seen in epithelial tissues, including the prostatic epithelium (THE GTEX CONSORTIUM et al., 2015; Tomlins et al., 2012). Out-of-context ERG activations via chromosomal translocations and fusion with different tissue-specific gene promoters trigger carcinogenesis in the organs of diverse developmental lineages. Prostate cancers (PCa) (Cancer Genome Atlas Research Network, 2015; Tomlins et al., 2005), Ewing sarcomas (Grünewald et al., 2018), or acute myeloid leukemia (AML) (Tsuzuki et al., 2011) are some of the exemplars of ERG-induced cancers. Identifications of ERG targets have primarily been based on genome-wide screening of its potential binding sites. One such study in PCa revealed that a TMPRSS2 promoter-driven ERG (Tomlins et al., 2005), which partners with HOXB13 and FOXA1 for its binding to its targets distributed throughout the genome; subsets of these targets show enrichment of Notch (N) signaling pathway members (Kron et al., 2017). In PCa, oncogenic ERG also upregulates Wnt ligand production (Wu et al., 2013), EZH2, a member of the Polycomb group (PcG) complex (Goel et al., 2021a; Yu et al., 2010), and DLX1, a homeobox transcription factor (Goel et al., 2021b). A hallmark of ERG-induced cancers, like PCa, is phenotypic plasticity—meaning changing morphological and functional characteristics *en route* to their metastatic progression [for review, see (Davies et al., 2018)]. These hallmarks of ERG-induced cancer, however, are not readily explained, despite the sheer number and diversities of ERG targets identified so far. Key challenges here are those of functional annotations of these ERG targets and deciphering their gene regulatory networks.

It has long been recognized that heterologous expression of mammalian transcription factors in *Drosophila* identify targets that display deep homology. Seminal reports on homeotic transformations in *Drosophila* following heterologous expression of mice Antennapedia (Antp)-like master transcription factor, Hox-2.2, (Malicki et al., 1990) or that of mammalian Pax6 master transcription factor-induced ectopic compound eyes (Halder et al., 1995), for instance, are exemplars of deep homology across distant phylogenies. Like Hox-2.2 or Pax6, ERG displays an ancient metazoan origin (Lautenberger et al., 1992), and its evolution is marked by the conservation of its DNA-binding ETS and PNT domains [for review, see (Hollenhorst et al., 2011)].

Here we reveal ERG oncoprotein-mediated repression of *Chip* in *Drosophila*. Further, reminiscent of that seen in PCa (Yu et al., 2010), ERG also binds to *E(z),* the *Drosophila* homolog of mammalian *EZH2*. ERG-induced E(z) represses *Chip* transcription via trimethylation of Lys27 of histone 3 of its promoter. *Chip* repression, in turn, disrupts the Chip-Tup, LIM-HD complex, alleviating repression of Notch-triggered Wg ligand synthesis in the posterior notum. Thus, a heterologous gain of ERG induces the development of Wg-mediated ectopic wing in the notum or drives tumor progression in *lgl* mutant notal cells. Our findings thus reveal that ERG represses a Chip-Tup, LIM-HD, complex that induces out-of-context Wg signaling triggering cell fate switches or carcinogenesis. Finally, we show that ERG-positive PCa cells display downregulation of *LDB1,* suggesting conservation of a crucial target of ERG repression identified from *Drosophila*. These results reveal a strategy to functionally identify the targets of human oncoprotein from *Drosophila* based on deep homology in their gene regulatory networks.

## RESULTS

### Human ERG oncoprotein induces ectopic dorsal appendages in *Drosophila*

*Drosophila* wing imaginal disc displays dorsal-ventral (D-V) and anterior-posterior (A-P) compartmentalization. A LIM-HD, Apterous (Ap), and a homeobox, Engrailed (En), transcription factor, respectively, marks its dorsal (D) and posterior (P) compartments [**Fig. 1A**, for a recent review, see (Wang and Dahmann, 2020)]. D-V and A-P boundaries double up as organizers of this appendage, secreting long-range Wg and Dpp morphogens, respectively [for a recent article, see (Zecca and Struhl, 2021)]. The presumptive adult wing (pouch domain) in the imaginal disc is marked by a POU-domain protein, Nubbin (Nub), which is a Wg target (Ng et al., 1996); it also marks the inner fold of the presumptive wing hinge (blue arrowhead, **Fig. 1B**).

**Fig. 1.**
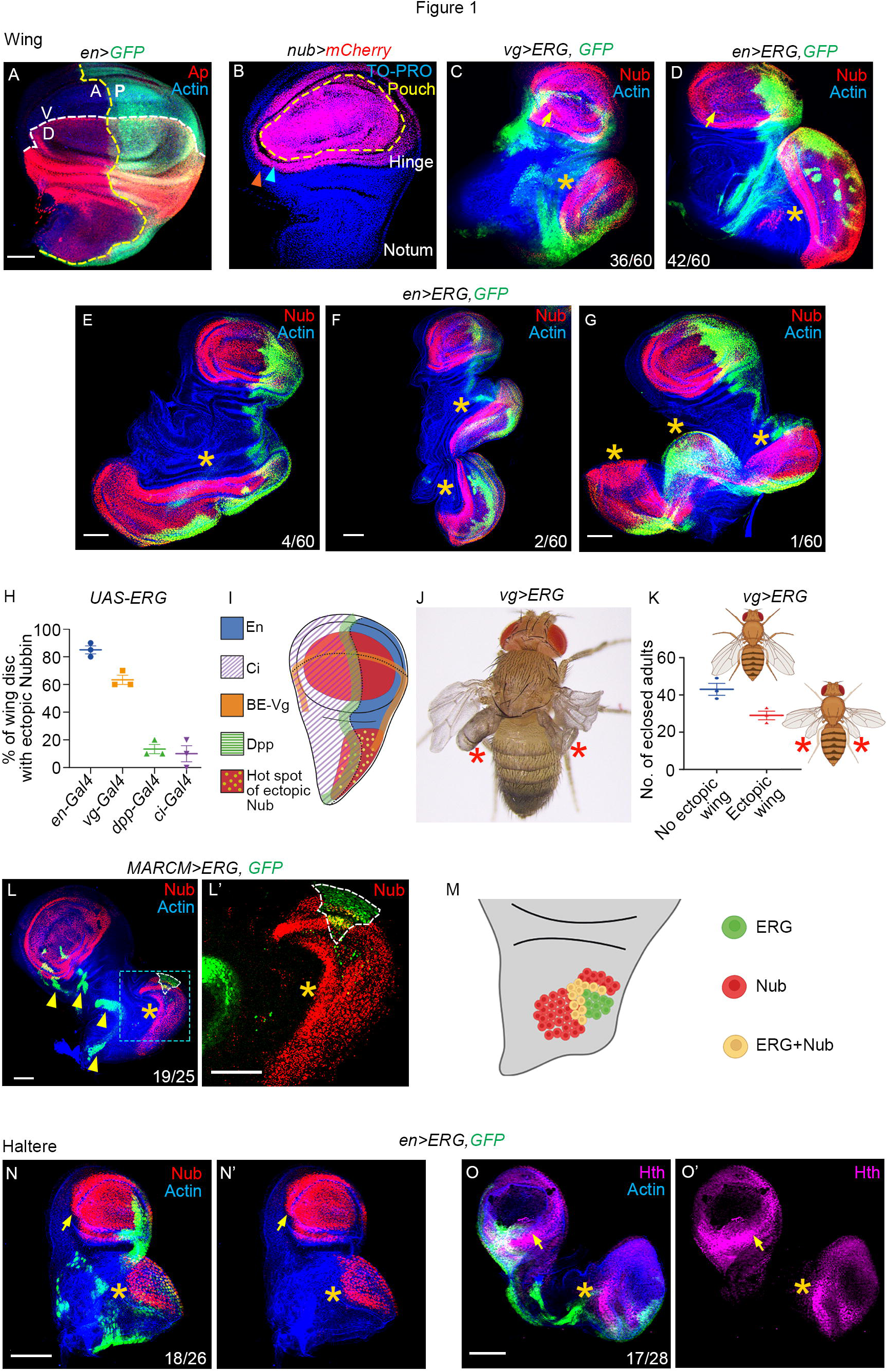
Heterologous ERG induces ectopic dorsal appendage primordia. (**A, B**) Apterous (red, Ap, A) and *en>GFP* (green, A), respectively, marks the dorsal (D) and posterior (P) compartments in the wing imaginal disc, whereas ventral (V) and anterior (A) compartments are their respective complementary domains. Yellow and white broken lines mark the A-P and D-V compartment boundaries, respectively (A). Nub (red, *nub>mCherry*, B) expression marks the presumptive adult wing (pouch, broken line, B) and the inner (blue arrowhead, B), but not the outer epithelial fold (orange arrowhead, B) of the hinge and notum. (**C, D**) ERG expression under *vg-Gal4* (*vg>ERG, GFP,* green, C) or *en-Gal4* (*en>ERG*, *GFP,* green, D) drivers induce ectopic Nub in the notum (star, C, D) reminiscent of that seen in the endogenous wing pouch domain (arrow, C, D). Note that yellow star here onwards marks the ectopic wing fate in the notum. (**E-G**) Examples of *en-Gal4>ERG* wing imaginal disc displaying an expanded (star, E) or two (star, F) or more than two (stars, G) ectopic Nub (red, E-G) expressing wing primordia in the notum. (**H, I**) Percent wing discs displaying ectopic Nub (wing pouch domain) expression in the posterior notum upon gain of ERG under *en-Gal4, vg-Gal4, dpp-Gal4,* or *ci-Gal4* driver (H). Schematic display of domains of these Gal4 expressions and their overlap with a hot spot of ectopic Nub induction in the posterior notum (I). (**J, K**) A *vg>ERG* adult displaying ectopic wings (stars, J). Quantification of ectopic adult wings (uni- or bi-lateral, K). (**L-M**) ERG-expressing MARCM clones (GFP, green, L). Its magnified view reveals non-cell-autonomous activation of Nub (star, red, L’) around the ERG clone (broken line, *GFP*, green, L’). Schematic representation of these results (M). (**N-O**) Gain of ERG in haltere imaginal discs induces ectopic Nub (star, *en>ERG*, N, N’) and Hth (star, purple, O, O’) reminiscent of their respective endogenous counterparts (arrows, L, L’, and O, O’). Scale bars-50µm; N= number of discs with desired marker or phenotype/ total wing discs observed

We first drove human ERG oncoprotein in larval wing imaginal disc under four different Gal4 drivers. These were *en-Gal4*, *vg-Gal* (boundary enhancer-BE)*, ci-Gal4,* and *dpp-Gal4* (see Supplemental Information, SI). Gain of ERG under *vg-Gal4* (star in **Fig. 1C**) or *en-Gal4* (stars in **Fig. 1D-G)** driver, for instance, induces ectopic Nub in the notum, reminiscent of its endogenous expression in the wing pouch. In extreme scenarios, albeit infrequent, the ectopic Nub-expressing wing primordium displayed growth far beyond its endogenous counterpart (star, **Fig. 1E**), while in others, induced two (stars, **Fig. 1F**) to three (stars, **Fig. 1G**) ectopic wing primordia in the notum. Frequencies of ERG-induced ectopic Nub expression in the notum progressively declined in the order of the Gal4 drivers used: *en-Gal4*, *vg-Gal4, dpp-Gal4,* and *ci-Gal4* (**Fig. 1H, Fig. S1A-C,** see SI). By plotting the domains of ectopic Nub gained under these Gal4 drivers, we could further trace a hot spot of its induction in the posterior compartment of the notum and straddling the A-P boundary (**Fig.1I**). Ectopic wing primordia induced by ERG expression displayed growth and patterning along the three developmental axes: A-P, D-V, and P-D (proximal-distal) (**Fig. S1D-F**). Only a fraction of the *vg>ERG* larvae (**Fig. 1J**) eclosed as adults displaying ectopic wings from the thorax (**Fig. 1J, K; Fig. S1G, G’**), while the rest showed larval/pupal lethality. Adult survival of only *vg>ERG* animals (**Fig. 1J**) appear to be linked to its selective expressions in dorsal appendages, like wing and haltere (see SI). ERG-expressing somatic clones, too, induced non-cell-autonomous expression of Nub (star, **Fig. 1L, L’**), but only when these were formed in the hot spot of posterior notum (see **Fig. 1I**), compared to those formed outside this territory (arrowheads, **Fig. 1L**). Therefore, these ERG-expressing clones in the notum displayed hallmarks of a morphogen-sending organizer [**Fig. 1M**; also see, (Bajpai and Sinha, 2020; Thompson et al., 2005).

In the haltere imaginal discs, a gain of ERG induced ectopic Nub- (**Fig. 1N, N’**) and Homothorax, Hth (**Fig. 1O, O’**), reminiscent of that seen in the wing imaginal discs. These outcomes are readily reconciled based on the shared developmental ground plans of these two dorsal appendage primordia (Weatherbee et al., 1998).

### ERG induces Wg morphogen-sending organizer in the posterior notum

During early, mid, to late third instar larval development, expression of Wg progressively elaborates the pouch, hinge, and notum domains of the wing imaginal disc [(**Fig. 2A**), for recent articles, see (Parker and Struhl, 2020; Zecca and Struhl, 2021)]. We noted an ectopic Wg expression in the *en>ERG* notum (stars, **Fig. 2B-D**), which appeared to be distinct from its wing counterpart till the late third larval instar (arrow, **Fig. 2B-D**). However, during an extended period of larval development, ectopic Wg expression in the notum (>6d AEL, broken yellow line, **Fig. 2E**) matched that of its endogenous counterpart (broken blue line, **Fig. 2E)** besides growing past the latter (see **Fig. 1E**). The endogenous wing primordium also displayed size reduction, presumably due to ERG-induced loss of Wg morphogen synthesis in its posterior wing compartment (blue arrowhead, **Fig. 2B’-D’**). We also noted that ERG-expressing clones selectively induced both cell-autonomous and non-cell-autonomous Wg in the posterior notum (star, **Fig. 2F, F’**), suggesting its organizer-like attribute (**Fig. 2G**).

**Fig. 2.**
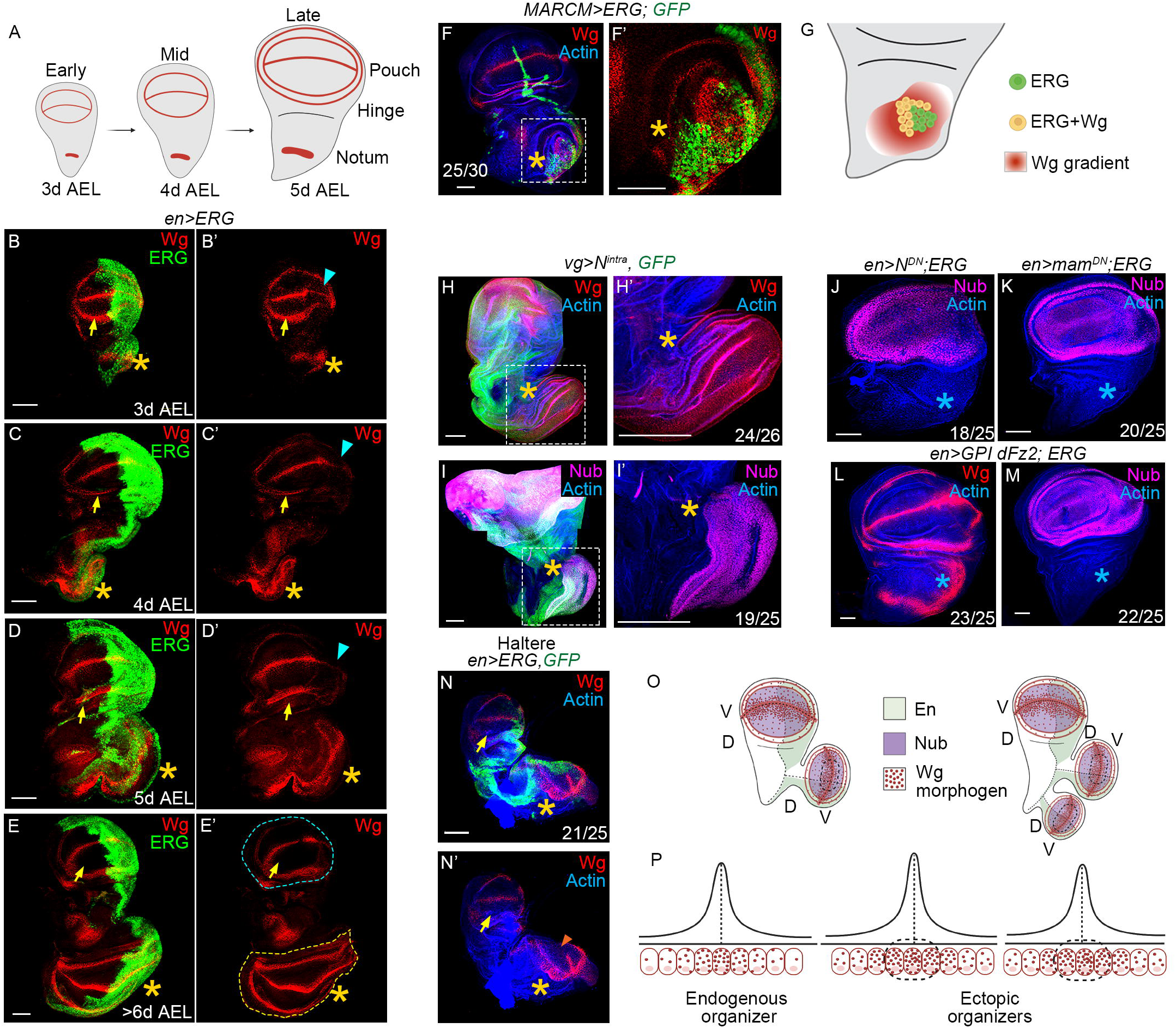
ERG-induces ectopic Wg morphogen-sending organizer of dorsal appendages. (**A**) Schematic display of Wg expression during third instar wing imaginal disc development. Larval age here and in subsequent images are shown in days after egg laying (d AEL). (**B-E**) Early (B), mid (C), late (D), and extended (E) third instar *en>ERG* wing discs display ectopic Wg expression (stars, B-E) reminiscent of the respective endogenous pattern (arrows, B-E). Comparison of sizes of fully grown ectopic (star) and endogenous (arrow) wing primordia (E, E’) are indicated by yellow and blue broken lines, respectively. Note the loss of endogenous Wg expression in the *en>ERG* (blue arrowheads, B’-D’) wing discs. (**F, G**) Mosaic wing epithelium of ERG-expressing clones (GFP, green, F) reveal ectopic and non-cell-autonomous gain of Wg (star, red, F’). Cartoon representation of these results (G). (**H-M**) Constitutive gain of Notch (*vg>N^intra^, GFP*, green, H, I) displays ectopic Wg (red, star, I) and Nub (purple, star, I) in the posterior notum; boxed areas in (H, I) are shown at higher magnification in (H’, I’). Co-expression of ERG with a dominant-negative form of N (blue star, *en>ERG; N^DN^*, J) or its downstream effector Mam (blue star, *en>ERG; mam^DN^*, K) extinguishes ectopic, ERG-induced, Nub gain in the notum. Co-expression of *GPI-dFz2* and *ERG* (*en>ERG; GPI-dFz2*) sequesters secreted Wg ligand (blue star, L) from the presumptive morphogen-sending ectopic organizer and arrests induction of ectopic Nub in the posterior notum (blue star, M). Note blue star here onwards marks the suppression of ectopic wing fate in the notum. (**N, N’**) *en>ERG*; *GFP* haltere imaginal disc displaying ectopic capitellum-like Wg (star, red, N, N’) like its endogenous counterpart (arrows, N, N’). The absence of Wg expression in the posterior D-V margin is a hallmark of haltere capitellum (orange arrowheads, N’). (**O, P**) Schematic interpretation of the results from (Figs. 1 and 2) as a fallout of induction of an ectopic D/V organizer(s) (O); thresholds of endogenous and ectopic Wg gradients are displayed as curves (P). Scale bar-50µm; N= number of discs with desired marker or phenotype/ total wing discs observed

The D-V organizer in the wing primordium displays an N-dependent Wg morphogen synthesis [see (Giraldez and Cohen, 2003)]. We also noted that gain of an activated N receptor (Struhl et al., 1993) under *vg-Gal4* (*vg>N^intra^*) displayed ectopic Wg (star, **Fig. 2H, H’**) and Nub (star, **Fig. 2I, I’**) in the posterior notum, reminiscent of those seen upon the gain of ERG (see Fig. 1C, D). Further, co-expression of ERG and a dominant-negative form of N, namely, N^DN^ (Rebay et al., 1993), or that of its downstream target, Mastermind, Mam^DN^ (Helms et al., 1999), extinguished ectopic gain of Nub in the posterior notum (blue star, **Fig. 2J, K**). Likewise, co-expression of ERG and a membrane-anchored receptor of Wg ligand, GPI-dFz2 (Cadigan et al., 1998)—which sequesters Wg (blue star, **Fig. 2L**)—also suppressed gain of the Wg target, Nub, in the notum (blue star, **Fig. 2M**). Finally, downregulation of *EGFR* signaling—which antagonizes Wg signaling (Szüts et al., 1997)—induced ectopic wing [(**Fig S2A-D**), also see (Baonza et al., 2000; Wang et al., 2000)]: that is, phenocopy gain of ERG (**Fig. 1**) or N (**Fig. 2**). These results, therefore, suggest that ERG antagonizes EGFR signaling in the notum, presumably via gain of Wg signaling [see (Baonza et al., 2000; Wang et al., 2000)]. ERG gain also induced ectopic Wg expression in the proximal haltere imaginal disc (star, **Fig. 2N, N’**) and induced Wg expression reminiscent of its endogenous capitellum primordium (arrow, **Fig. 2N, N’**).

Therefore, a heterologous gain of ERG oncoprotein in the adult thorax primordia induces the development of ectopic dorsal appendages by setting up one or more than one N-dependent Wg morphogen-sending organizers *de novo* (**Fig. 2O, P**).

### ERG disrupts a LIM-HD, Chip-Tup, complex by repressing *Chip*

LIM-HD protein complexes maintain developmental domain-specific cellular signaling and cell fate in *Drosophila* imaginal discs (Milán and Cohen, 1999; de Navascués and Modolell, 2007; Roignant et al., 2010). For instance, Chip-Tup (Biryukova and Heitzler, 2005; de Navascués and Modolell, 2007) and Chip-Ap (Fernández-Fúnez et al., 1998; van Meyel et al., 1999; Milán and Cohen, 1999) are, respectively, active in the proximal (notum) and distal (wing) domains of the wing imaginal disc.

Given that the Chip-Tup LIM-HD complex regulates notum development (Biryukova and Heitzler, 2005; de Navascués and Modolell, 2007), we examined the fallouts of individual losses of Chip and Tup in this domain. Gain of a dominant-negative form of Chip, Chip^Δoid^ (Torigoi et al., 2000) in somatic clones (**Fig. 3A**), or via ubiquitous expression (*en>Chip*^Δ^*^oid^*, **Fig. 3B**) induced Nub expression in the notum. Further, *vg-Gal4>Chip*^Δ^*^oid^* adult displayed an ectopic wing in the thorax (**Fig. 3C**). Loss of Chip, therefore, phenocopies ERG gain (Figs. 1, 2), suggesting the possibility of its repressive binding on the promoter of the former. We noticed two putative ERG-binding sites (EBS) within ∼2 kb upstream of the *Chip* promoter (EBS-1 and EBS-2, **Fig 3D**). These EBSs were also recovered from chromatin immuno-precipitates (ChIP) of *en>ERG* wing imaginal discs using ERG antibody (**Fig. 3E, F**). Further, co-expression of Chip and ERG extinguished ectopic Nub (star, **Fig. 3G**) or Wg (star, **Fig. 3H**), overriding the presumed repressive binding of ERG on the *Chip* promoter. Moreover, in rare adults emerging from *vg>Chip; ERG* larvae also failed to display the characteristic ectopic wing from the thorax (star, **Fig. 3I**), unlike their counterparts with individual gain of either ERG (Fig. 1G) or loss of Chip (Fig. 3C).

**Fig. 3.**
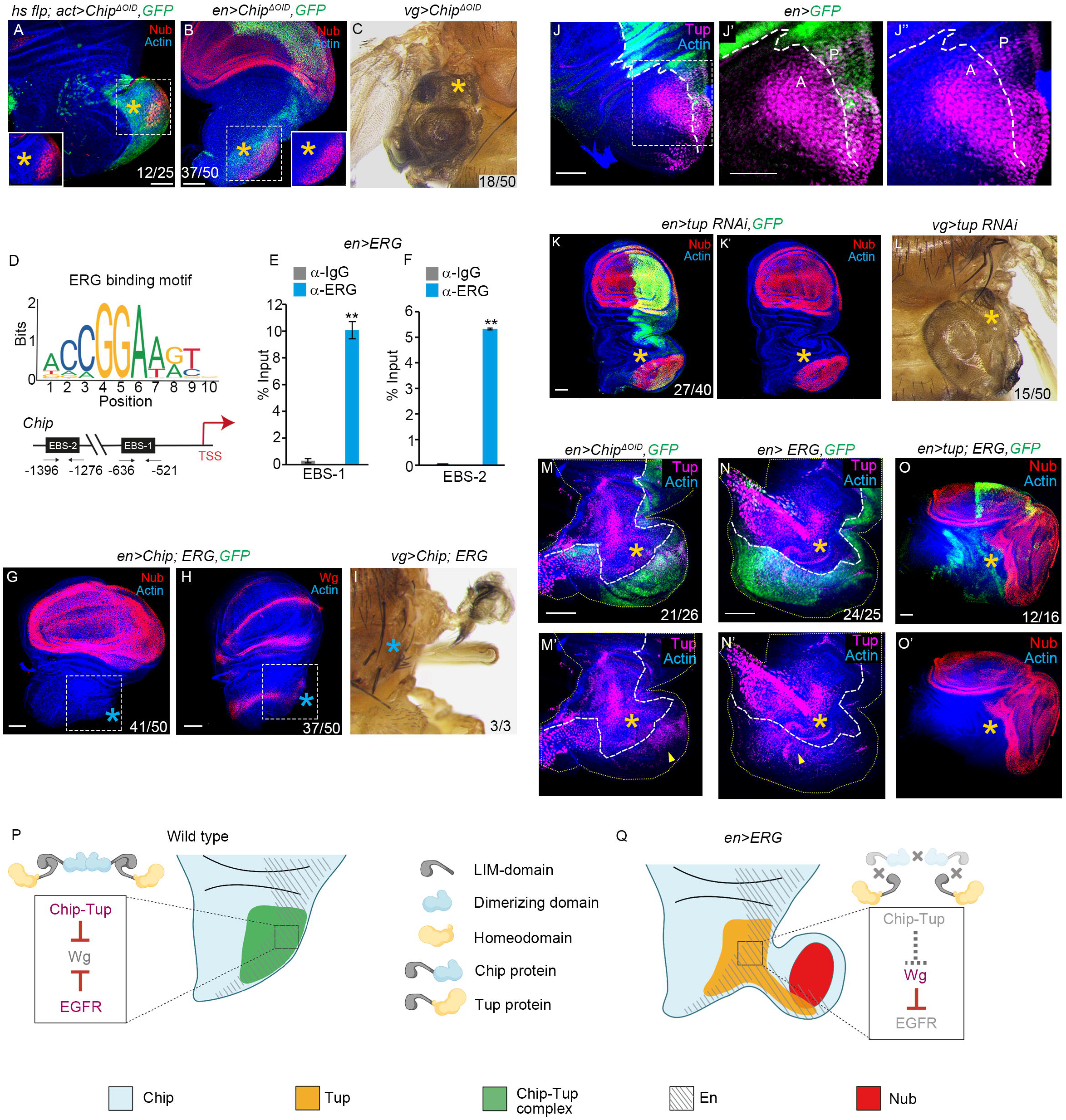
Heterologous ERG abrogates the Chip-Tup complex by targeting *Chip* expression. (**A**-**C**) Gain of a dominant-negative form of Chip (Chip^Δoid^) in a somatic clone (GFP, A) or under an *en-Gal4* driver (*en> Chip*^Δ^*^oid^; GFP*, green, B) induce Nub (star, A, B) in posterior notum (insets display only the Nub channel). *vg>Chip*^Δ^*^oid^* adult displaying ectopic wing in the adult thorax (star, C). (**D-F**) Consensus ERG binding motif (top, D) and a -2kb sequence upstream of *Chip* promoter displaying two ERG binding sites (EBS-1 and EBS-2, bottom, D). Enrichment of ERG binding on EBS1 and EBS2 in ChIP-qPCR from *en>ERG* wing imaginal discs using anti-ERG antibody (E, F). \*\**p*<0.001 (**G-I**) Co-expression of Chip and ERG (*en>Chip, ERG)* suppresses induction of ectopic Nub (blue star, G) or Wg (blue star, H) in the notum or the development of ectopic wing from the adult thorax (blue star, *vg>Chip; ERG,* I). (**J-J”)** Expression of Tup (red) in the notum of third instar wing imaginal disc (J); broken line marks the A-P compartment boundary (*en>GFP;* green, J). Boxed area of (J) is shown at higher magnification in (J’, J’’) (**K-L**) Knockdown of *tup* in wing imaginal disc (*en>tup-RNAi*, GFP, green, K) induces ectopic Nub (star, K, K’) while *vg>tup-RNAi* adults display ectopic wing in the thorax (star, L). (**M-N**) Loss of *Chip* (green, *en> Chip*^Δ*oid*^, M) or gain of ERG (green, *en> ERG*, N) in wing imaginal discs fail to suppress Tup (red, M’, N’) in the notum. (**O**) Co-expression of *tup* and *ERG* fails to suppress ectopic Nub (star, *en>tup; ERG*, O, O’; for comparison, see G). (**P, Q**) Schematic interpretation of results displayed in Fig. 3 and Fig. S2. Scale bar-50µm; N= number of discs with desired marker or phenotype/ total wing discs observed

The expression of Tup—a partner in the Chip-Tup tetrameric complex—is restricted to the posterior notum in the third instar wing imaginal disc [**Fig. 3J**, see (Biryukova and Heitzler, 2005), while expression of *Chip* is ubiquitous throughout the wing imaginal discs (Morcillo et al., 1997). A notum-specific expression of Tup may thus underlie the fallout of the loss of Chip (Fig. 3A-C) or gain of ERG (Figs. 1-2). We noticed that knockdown of *tup* induced ectopic Nub (*en>tup RNAi*, **Fig. 3K, K’**) and wing from the notum (**Fig. 3L**), phenocopying Chip loss (Fig. 3A-C), and ERG gain (Fig. 1, 2). These results reveal that Chip-Tup complex antagonizes Wg ligand synthesis in the notum; thus, loss of any of these two LIM-HD partners results in comparable fallout in this domain due to disruption of the complex. We further noticed that neither loss of Chip (**Fig. 3M**) nor gain of ERG (**Fig. 3N**) extinguished Tup expression in the notum, revealing thereby that ERG-induced Nub expression is not causally linked to loss of Tup *per se*. Moreover, unlike that of Chip (see Fig. 3G, H), gain of Tup failed to reverse ERG-mediated ectopic Nub expression (*en> tup; ERG,* **Fig 3O, O’**).

Together, these results (Figs. 2-3) show that Chip-Tup complex- and EGFR-mediated suppression of Wg ligand synthesis in the posterior notum (**Fig. 3P**) is destabilized by ERG-induced repression of *Chip* (**Fig. 3Q**). Further, although Tup *per se* is not repressed by a gain of ERG (**Fig. 3Q**), it is rendered non-functional in the absence of its partner, Chip.

In contrast to the notum, in the wing proper, another LIM-HD complex, Chip-Ap, promotes wing development via an N-dependent Wg ligand synthesis (Cohen et al., 1992; van Meyel et al., 1999; Milán and Cohen, 1999). We noticed that ERG gain repressed N-Wg signaling cascade in the wing proper, even while it did not directly target Ap (**Fig. S3A-G**). However, these opposing fallouts of ERG gain in the wing and notum primordia are readily reconciled by ERG-mediated repression of *Chip* (compare **Fig S3H, I** with **Fig. 3P, Q**).

### ERG-induced upregulated E(z) epigenetically silences *Chip*

Direct binding of ERG on *EZH2* (Kleer et al., 2003; Koyanagi et al., 2005; Varambally et al., 2002; Yu et al., 2010), a homolog of *Drosophila E(z)* (Laible et al., 1997), upregulates expression of the latter in different cancers. In *Drosophila,* too, we noticed a putative ERG-binding site (EBS) between -272 to -183 bp upstream in the *E(z)* promoter (**Fig. 4A**), which was further confirmed in ChIP of *en>ERG* (**Fig. 4B)** wing imaginal discs.

**Fig. 4.**
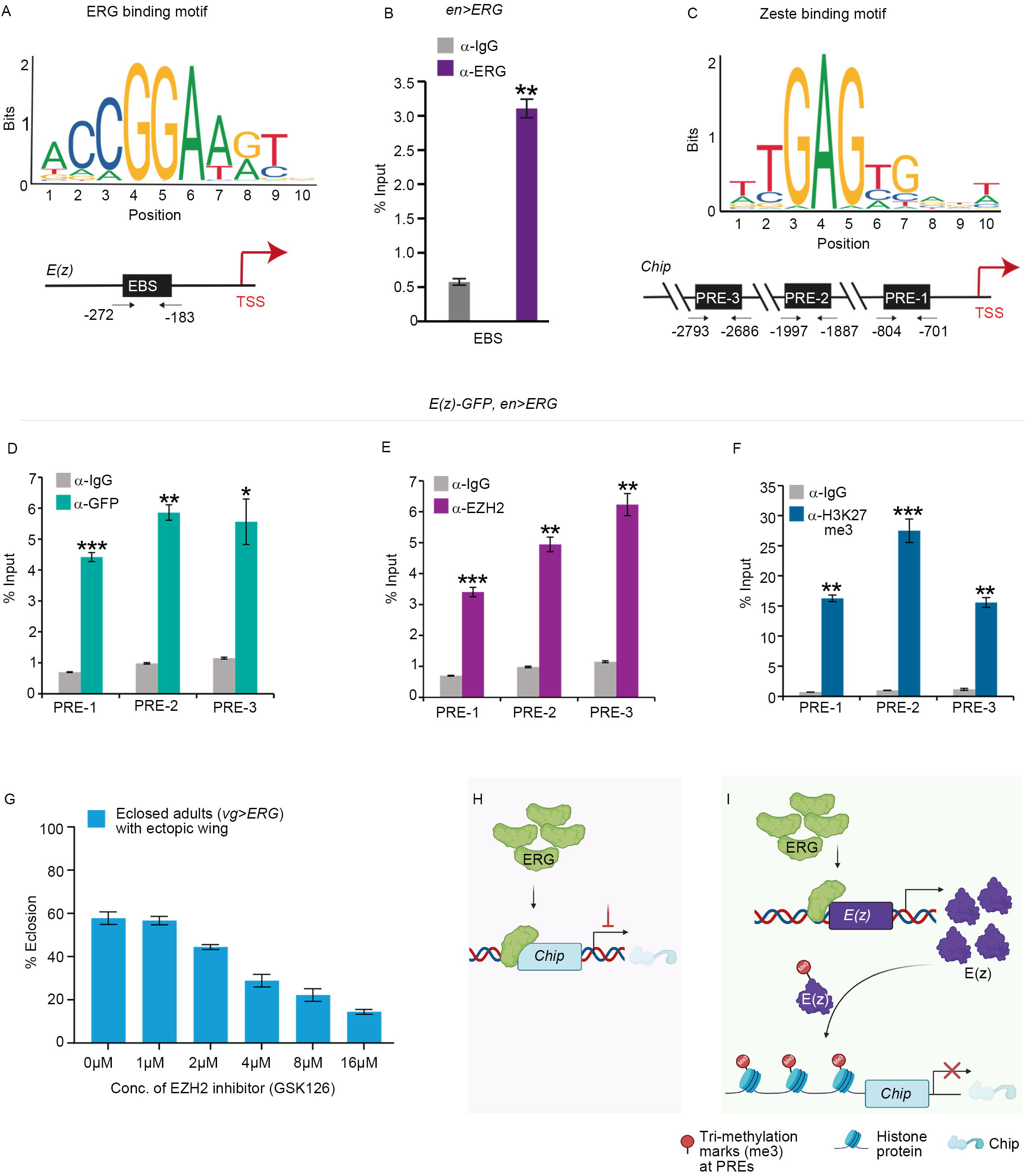
ERG binds to both *Chip* and E(z), and the latter epigenetically silences the former. (**A, B**) Consensus ERG binding sequence motif (top, A), within a -2 kb of *E(z)* (EBS, bottom, A) from TSS (transcription start site). Enrichment of this EBS in ChIP-qPCR from *en>ERG* wing imaginal discs (B). (**C-F**) Consensus Zeste binding sequence motif (top, C) within a -3 kb *Chip* TSS displays three Polycomb response elements: PREs (PRE1, PRE2, and PRE3 (bottom panel, C). Enrichment of these PREs in ChIP-qPCR from *E(z)-GFP*; *en >ERG* wing imaginal discs following pulled down with anti-GFP (D), mammalian anti-EZH2 (E), or anti-H3K27me3 (F). \**p*<0.01, \*\**p*<0.001, \*\*\**p*<0.0001 (**G**) Larva fed on increasing concentrations of EZH2 inhibitor, GSK126, progressively displayed suppression of ERG-induced (*vg>ERG*) ectopic wing development in eclosed adults. (**H, I**) Schema displaying ERG-mediated regulation of *Chip* by its direct binding (H), and via E(z) (I).

E(z) is a transcriptional repressor, a core component of the Polycomb repressive complex 2, PRC2, which binds to <u>P</u>olycomb <u>R</u>esponse <u>E</u>lements (PREs) of its targets [see (Schuettengruber et al., 2007)]. We noticed three PREs within ∼3 kb upstream of the *Chip* transcription start site, TSS (**Fig. 4C**). These were enriched in the ChIP of ERG-expressing wing imaginal discs—which also carried an endogenous E(z)-GFP fusion protein as well (Kudron et al., 2018); *E(z)-GFP, en>ERG*)—upon pull-down by anti-GFP (**Fig. 4D**) or anti-EZH2 (*E(z)-GFP, en>ERG*, **Fig. 4E**) antibodies. These PREs of *Chip* also displayed the trimethylation of histone3K27, H3K27me3 (**Fig. 4F**), a hallmark of E(z)-mediated epigenetic silencing [see (Schuettengruber et al., 2007)]. We further asked if an E(z)-mediated silencing of *Chip* upon ERG gain could be reversed pharmacologically (Kim et al., 2018). To test this possibility, we fed *vg>ERG* larvae on food supplemented with a well-characterized inhibitor of EZH2, GSK126 (Kim et al., 2018). Indeed, *vg>ERG* adults eclosing from larvae fed on GSK126-supplemented food displayed a progressive and concentration-dependent reversal of ectopic wing development (**Fig. 4G).**

Thus, while ERG binding on the *Chip* promoter is likely to be repressive (**Fig. 4H**), an ERG-induced and E(z)-mediated H3K27me3 on *Chip* PREs provides further definitive evidence of its epigenetic silencing (**Fig. 4I**). These outcomes (**Fig.1-4**) are also reminiscent of phenotypic plasticity seen in cancer progression, especially those induced by ERG gain (Yu et al., 2010).

### In the posterior notum, ERG-induced disruption of the Chip-Tup complex triggers tumor cooperation

*Drosophila* displays a well-known two-hit paradigm of carcinogenesis (Brumby and Richardson, 2005; Khan et al., 2013; Pagliarini and Xu, 2003). Indeed, *lgl* clones with a simultaneous ERG gain (*lgl; ERG*) displayed neoplastic transformation in the posterior notum (broken line, **Fig 5A, A’**), while in the rest of the wing imaginal disc, these often failed to transform (arrowheads, **Fig. 5A**). In the posterior notum, *lgl; ERG* clones also tend to induce Nub non-cell autonomously with accompanying overgrowth (star, **Fig. 5A**), reminiscent of their ERG-expressing counterparts in the posterior notum (see Fig. 1L, L’). Further, in unfixed mosaic wing epithelium, secreted Wg (Strigini and Cohen, 2000) was seen in both transformed *lgl; ERG* clone in the notum (broken line, **Fig. 5B, B’’**) and in the neighboring hyperplastic domains marked by excessive epithelial folds (arrow, **Fig. 5B’, B”’**), reminiscent of morphogen-sending neoplastic clones (Bairzin et al., 2020; Bajpai and Sinha, 2020). Cell neighbors of such Wg ligand-sending *lgl; ERG* clones also displayed excessive cell proliferation (arrow, PH3, red, **Fig. 5C, C’**).

**Fig. 5.**
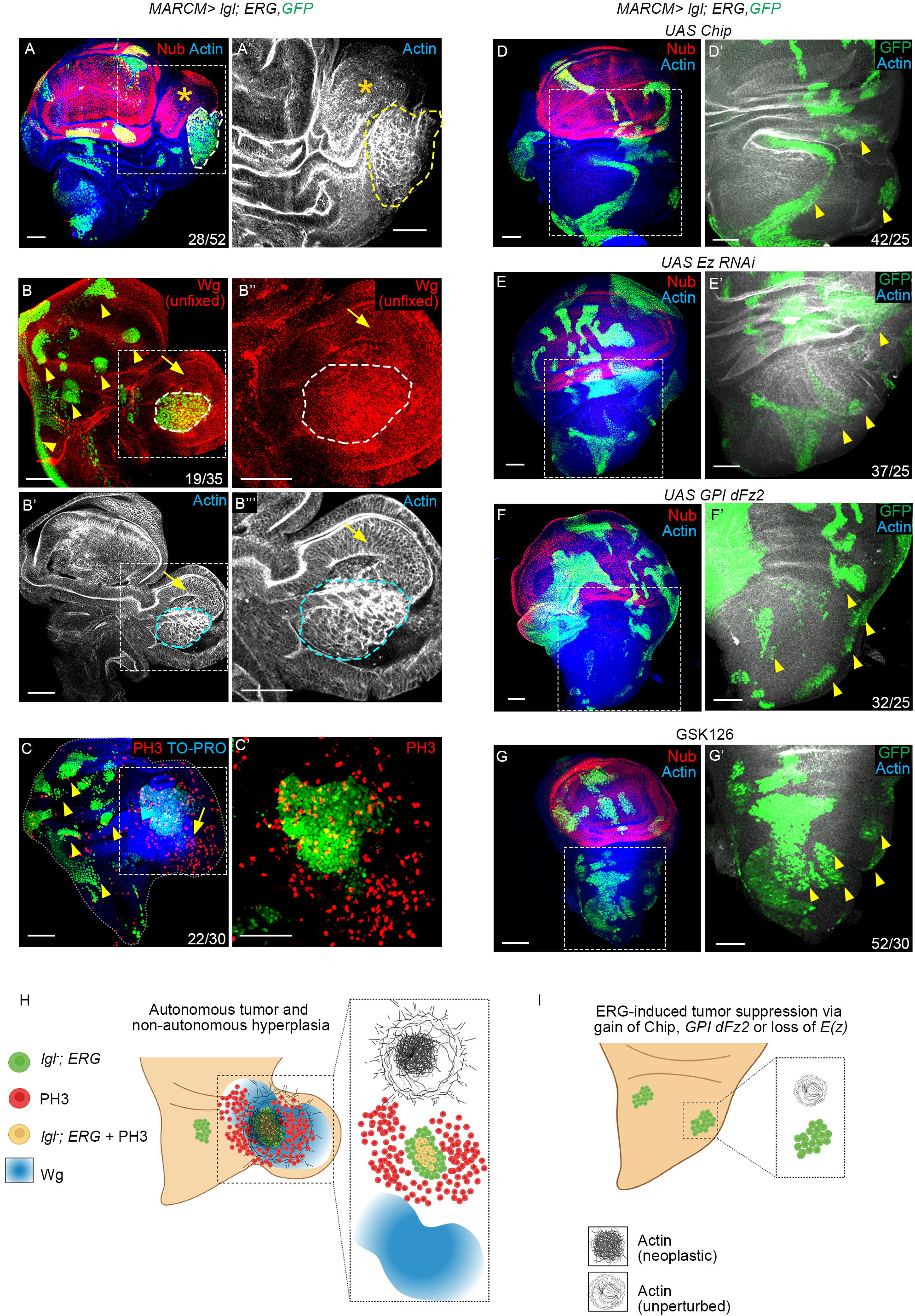
ERG-induced *lgl* tumorigenesis in posterior notum via Wg morphogen signaling. (**A, A’**) An ERG-expressing *lgl* clone (GFP, green, A) in the posterior notum; boxed area in (A) is shown at higher magnification in (A’). Note the ectopic, non-cell-autonomous gain of Nub and hyperplasia in notal cells abutting the clone (star. A. A’), while neoplastic transformation remains autonomous (broken line, A, A’). (**B-B’”**) An unfixed mosaic wing imaginal disc epithelium displaying ERG-expressing *lgl* clones (GFP, green) and secreted Wg (red). Boxed areas in (B, B’) are shown at higher magnification in the right panels (B’’, B”’). Note autonomous neoplasia in the clone (broken line) while the surrounding epithelium display secreted Wg (arrow, B’) with accompanying hyperplasia marked by their high epithelial folding (arrow, B”’). Clones in other domains are not transformed (yellow arrowheads, B). (**C, C’**) A mosaic wing imaginal disc epithelium displaying *lgl; ERG,* clone (green), and stained for a cell proliferation marker phospho-histone, PH3 (red). The boxed area is shown at higher magnification (red, C’), revealing over proliferation of cells surrounding the *lgl; ERG* clone in the posterior notum (arrow, green, C’) unlike the rest (yellow arrowhead, C). (**D-G**) *lgl; ERG* clones in mosaic wing imaginal disc epithelia with simultaneous gain of Chip (D), knockdown of *E(z)* (E), expression of *GPI-dFz2* (F), or dissected from larvae fed on EZH2 inhibitor, GSK126 (25mg/ml) (G). Note that none of these mosaic epithelia display transformed clones (actin, blue arrowheads, D’-G’) or induce ectopic Nub (red, D-G). (**H, I**) Cartoon interpretations of selective neoplastic transformations in posterior notum (H) and their suppression (I). Scale bar-50µm; In each figure, N= the number of clones displaying the relevant phenotype out of the total number of wing discs examined.

Finally, we note that Chip gain (**Fig. 5D, D’**), *E(z)* knockdown (**Fig. 5E, E’**), or Wg sequestration (**Fig. 5F, F’**) in *lgl; ERG* clones arrested their neoplastic transformation. These clones also failed to display Wg signaling or activate its target, Nub (**Fig. 5D-G**, for comparison, see Fig. 5A). Comparable results were obtained upon feeding host larvae bearing *lgl; ERG* mosaic imaginal discs on an E(z) inhibitor, GSK126 (**Fig. 5G, G’**, also see Fig. 4G). Thus, ERG-induced tumors in *Drosophila* appear to recapitulate phenotypic plasticity in a select spatial context (**Fig. 5H, I**).

### In the wing hinge domain, *lgl, ERG* clones transform sans *de novo* Wg ligand synthesis

N signaling activates Wg ligand synthesis in the D/V organizer of the wing imaginal discs (Zecca and Struhl, 2021). Ectopic gain of an activated N receptor, N^intra^ (Struhl et al., 1993) in *lgl* clones recapitulates these hallmarks of Wg ligand synthesis and secretion: autonomous and non-cell-autonomous induction of Nub (*lgl; N^intra^* **Fig. S4).** *lgl; ERG* clones in the hinge domain of wing imaginal discs, however, neither displayed expansion of its Nub-expressing cells within the clone boundary nor induced its non-cell-autonomous expression (**Fig. 6A, B** for comparison, see Fig. 5A, B). These features of *lgl; ERG* clones in the hinge domains, unlike their counterparts in posterior notum (See Fig. 5A, B), reveal that transformation of these clones does not entail *de novo* Wg synthesis.

**Fig. 6.**
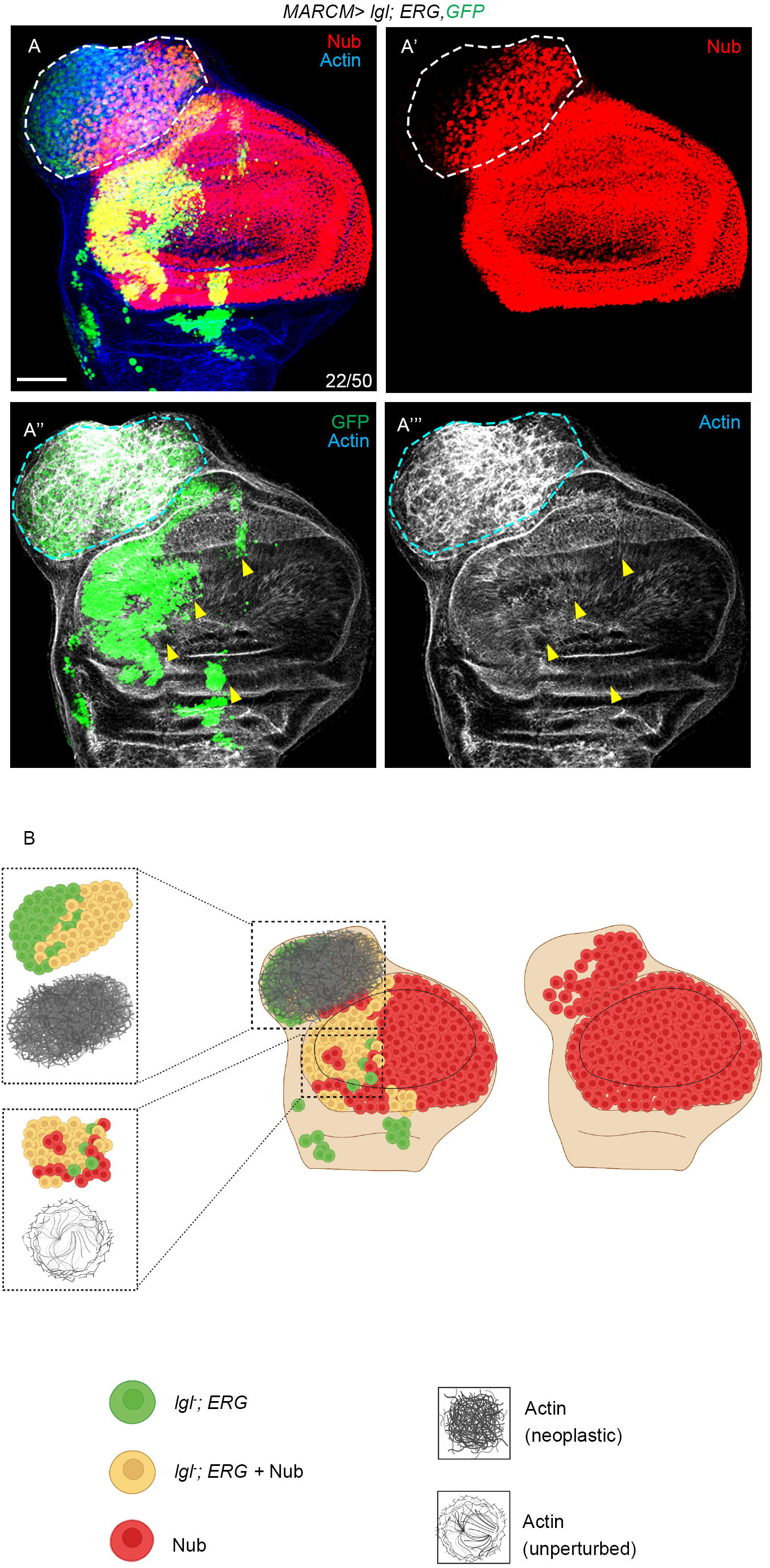
In the hinge domain, *lgl; ERG* clones do not induce Wg morphogen synthesis. (A) An ERG-expressing large *lgl* clone (GFP, green, A) in the hinge domain of a mosaic wing imaginal disc. This clone displays neoplastic transformation (actin, broken line A”’); however, not all cells of the clones do not display expression of a Wg target, Nub (red, white broken line, A’). *lgl; ERG* in the wing pouch does not display neoplastic transformation (yellow arrowheads). (B) Schematic interpretation of results displayed in (A-A’”). Scale bar-50µm; N= number of clones displaying the relevant phenotypic marker out the total test clones formed in the hinge

### Conservation of ERG-induced *Chip/LDB1* repression in prostate cancer cells

We noticed that the protein-protein interactions (PPI) map of *Drosophila,* centered on the Chip-Tup complex, was comparable with its mammalian counterparts [**Fig. S5A** also see (Te Velthuis et al., 2007)], suggesting an ancient origin of this complex. We further asked if ERG targets *LDB,* the mammalian homolog of *Drosophila Chip*. Unlike *LDB1* and *LDB2*, the two isoforms of mammalian *LDB* (Matthews and Visvader, 2003), ERG expression is not seen in the healthy prostatic epithelium (**Fig. S5B-G**). Therefore, ERG likely targets these isoforms of *LDB*s in cancers, such as PCa. We chose to test this hypothesis in ERG-positive or - negative PCa cell lines. Analysis of publicly available ChIP-Seq data of ERG-positive VCaP cell line [GSE28950, (Chng et al., 2012)] revealed binding peaks of ERG on the *LDB1* promoter (**Fig. 7A**) but not on that of *LDB2* (**Fig. 7B**). This peak was no longer seen in ChIP-Seq data of the VCaP cell line that displays knockdown of *ERG* [**Fig. 7C**, (GSE110655), see SI]. Thus, between the two *LDB* isoforms, only *LDB1* displayed binding of ERG, which was further confirmed by the presence of a putative ERG binding site (EBS) in its promoter (**Fig. 7D**). We could further detect its enrichment through chromatin immunoprecipitation from ERG-positive VCaP cells (ERG*_LDB1* **Fig. 7E, F**), suggesting that the binding of ERG on *LDB1* promoter could repress the latter. In agreement, upon knockdown of *ERG* in the VCaP cell line, we noted elevated *LDB1* expression (**Fig. 7G**). Conversely, we also noticed that an ERG-negative benign prostate cell line, RWPE, displays downregulation of *LDB1* upon *ERG* transfection (**Fig. 7H**, GSE86232, see SI), while its knockdown in ERG-positive VCaP cells is marked by upregulation of *LDB1* (**Fig. 7I**, GSE110656, see SI). Finally, we also noticed an *ERG* knockdown-mediated downregulation of *EZH2* expression (**Fig. 7J**, GSE110656, see SI) in VCaP cells. These results are consistent with the *EZH2* being a transcriptional target of ERG mammalian cancers (Chng et al., 2012; Yu et al., 2010) and also in *Drosophila* (Fig. 4).

**Fig. 7.**
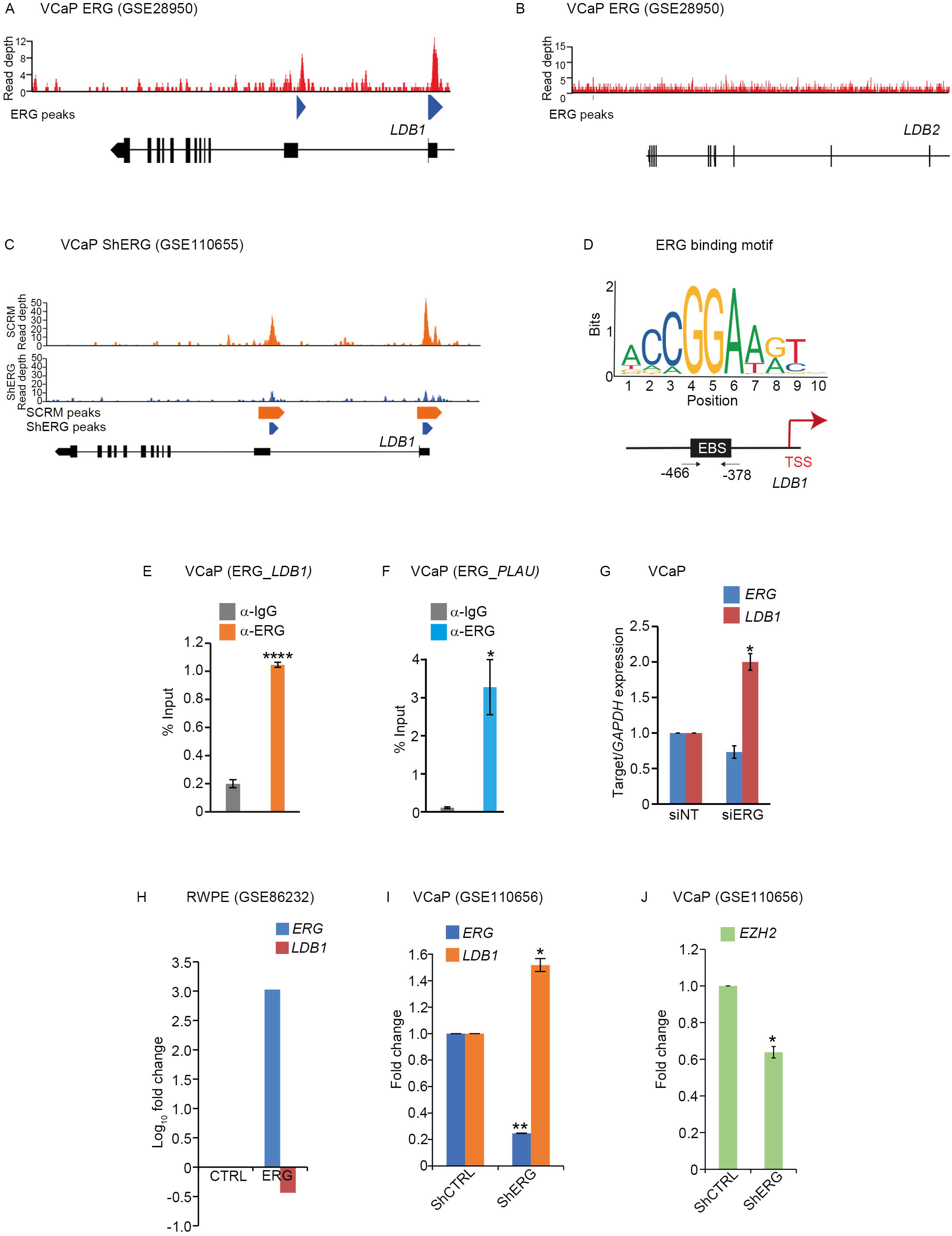
ERG positive VCaP PCa cell line display suppression of *LDB1*, a homolog of the *Drosophila Chip*. (**A, B**) ChIP-seq dataset of an ERG-positive VCaP cell line (GSE28950) displays strong binding peaks (blue arrowhead MACS, Model-based Analysis of ChIP-seq) of ERG on *LDB1* (A), which are missing in *LDB2* (B). (**C**) ChIP-seq dataset of an ERG-positive VCaP cell line (GSE110655) displays loss of ERG binding peaks on *LDB1* upon knockdown of ERG (blue, bottom panel) compared to control (orange, top panel). Note orange arrowheads marks the ERG-binding MACS peak in control while reduced blue arrowheads display loss of ERG-occupancy on *LDB1* upon ERG knockdown (**D**) *LDB1* promoter displays a binding site for ERG (EBS, D). (**E**, **F**) Enrichment ERG at the predicted EBS in ChIP-qPCR of *LDB1* promoter (E); a known ERG target, *PLAU (Plasminogen activator urokinase)*, served as an internal control (F). (**G**) Quantification of *LDB1* mRNA by qPCR following knockdown of *ERG* in VCaP cell line. (**H**) Gene expression analysis in benign prostate cell line (RWPE) displaying ERG over-expression (GSE86232) reveal downregulation of *LDB1*. (**I, J**) Gene expression analysis in ERG-positive VCaP cell line displaying ERG knockdown (GSE110656) reveals upregulated *LDB1* (\**p*<0.05, I) and downregulated *EZH2* (\*\**p*<0.005, J).

Together, these results reveal the functional conservation of ERG targets in human, which are identified from *Drosophila*.

## Discussion

Genetic tractability apart, *Drosophila* offers the advantages of deep homology of its essential patterning mechanisms with mammals (Shubin et al., 2009). Here we have uncovered a single causal underpinning—an out-of-context Wg ligand synthesis in the posterior notum of *Drosophila*—upon two independent genetic perturbations: namely, a heterologous gain of human ERG oncoprotein and downregulation of an endogenous gene, *Chip*, that codes for a LIM-domain binding protein. These genetic manipulations disrupt the Chip-Tup, LIM-HD complex, a repressor of Wg ligand synthesis in the posterior nota leading to the development of ectopic dorsal appendages in *Drosophila*. ERG binds to the promoters of both *Chip* and *E(z),* the latter finding being reminiscent of that seen in mammalian cancers (Chng et al., 2012; Yu et al., 2010). Epigenetic silencing of *Chip* via E(z) mediated trimethylation of H3 K27 of its promoter, therefore, underpins disruption of Chip-Tup complex. We also noted gain of *N* or *Wnt* signaling upon ERG gain in *Drosophila*, reminiscent of that seen in cancers (Kron et al., 2017; Tsuzuki et al., 2011; Wu et al., 2013). Our implication of the LIM-HD complex in carcinogenesis in *Drosophila* also resonates with the known roles of LIM-HD transcription complexes as negative regulators of Wnt signaling (Renko et al., 2019). LIM-HD complexes are deployed to recruit lineage-specific DNA binding proteins and co-activators (Fiedler et al., 2015; Renko et al., 2019). Targeting this complex by ERG readily reconciles its implication in cancers in multiple tissue lineages [see (Adamo and Ladomery, 2016)].

That ERG could be a transcriptional repressor of *Chip/LDB1* was not predictable from the large body of literature directed at the identification of ERG targets from different mammalian cancers [for reviews, see (Hollenhorst et al., 2011; Hsing et al., 2020; Lin et al., 2017; Qian et al., 2022)]. For instance, although we could find ERG binding sites (EBS) on *LDB1* from a ChIP-seq study reported previously [(Chng et al., 2012), see SI], its relevance in cancer progression was not evident in the absence of its functional annotation. Driving a human ERG in *Drosophila,* on the other hand, offers an unbiased approach to identify ERG-regulated target(s) *in vivo* based on a discernible phenotype, *à la* reverse genetics. The oncogenic relevance of this approach is further underscored by our display of ERG-mediated downregulation of *Chip/LDB1* in PCa cells. Previously described strategies in modeling cancers in *Drosophila* entailed driving fly orthologs of human oncogenes. Ectopic gain of *dRet^MEN2^* (Read et al., 2005) or different variants of *dRas^v12^* (Bangi et al., 2016, 2019; Brumby and Richardson, 2005; Pagliarini and Xu, 2003), for instance, are two exemplars of mimicking cellular pathogenesis of thyroid and colorectal cancer, respectively. The expression of human oncogenes/tumor suppressors like p53 (Yamaguchi et al., 1999) or RAS-kinases (Das et al., 2021) presents yet another approach to modeling cancer in *Drosophila*. The strategy shown in this study is likely to be most relevant in identifying oncoprotein targets and gene regulatory networks of oncogenically activated selector genes, including homeotic selectors, which are abundantly implicated in cancers [see (Jonkers et al., 2020; Shah and Sukumar, 2010)].

A crucial advantage of the phenotypic fallouts of oncoprotein gain in *Drosophila* is its insightful extension to the mammalian system. For instance, reactivation of Wg signaling in the notum is also seen upon loss of Osa (Collins and Treisman, 2000) or subunits of BAP (Brm-associated protein) (Song et al., 2017) chromatin remodeling complexes in *Drosophila*, which are conserved in mammals [for review, see (Kadoch and Crabtree, 2015)]. Furthermore, the binding of ERG to the BAP/BAF chromatin remodeling complex in PCa (Sandoval et al., 2018) now reconciles, given the results presented here, with an emergent ERG-targeted gene regulatory network. By extension, it will also be interesting to examine if tumor cooperation by *Drosophila* ortholog of human ERG, such as Ets21C (Külshammer et al., 2015; Mundorf et al., 2019; Toggweiler et al., 2016), entails its targeting the Chip-Tup complex.

ERG-driven *lgl* carcinogenesis revealed here is reminiscent of cell fate switches envisaged in a Waddington landscape (Berdasco and Esteller, 2010; Flavahan et al., 2017). Suppression of Chip-Tup complex represents one such notum-to-wing lineage switch, which turns oncogenic upon a second hit, namely, loss of Lgl (**Fig. 8**). However, only a select group of cells, namely, the cells of the posterior notum, displays this attribute marked by Wg ligand synthesis, which qualifies as cancer cells of origin (Khan et al., 2013; Visvader, 2011) in ERG-induced tumors in the posterior notum. In other words, spatial limits of expression of Chip-Tup-like LIM-HD complexes could be the determinants of tumor hot or cold spots [see (Tamori and Deng, 2017)]. Conversely, in the wing hinge domain, where Chip-Tup, LIM-HD complex is not formed (Milán and Cohen, 1999; Milán et al., 1998), *lgl; ERG* clones did not trigger Wg ligand synthesis and yet displayed neoplastic transformation, revealing distinct attributes of cancer cells of origin in the hinge. These findings are consistent with the different developmental outputs of spatial context-specific expression of LIM-HD complexes and their partnership with locally expressed transcription factors [see (Bhati et al., 2008; Matthews and Visvader, 2003)]. By extension, we speculate that intra-tumoral and spatial heterogeneity in ERG-driven cancers (Haffner et al., 2021; Mann et al., 2016; Minner et al., 2013) could be linked to cell type-specific recruitment LIM-HD binding partners. Indeed, LDB1-mediated Wnt signaling appears distinct in proximal and distal colorectal tumors (García et al., 2016).

**Fig. 8.**
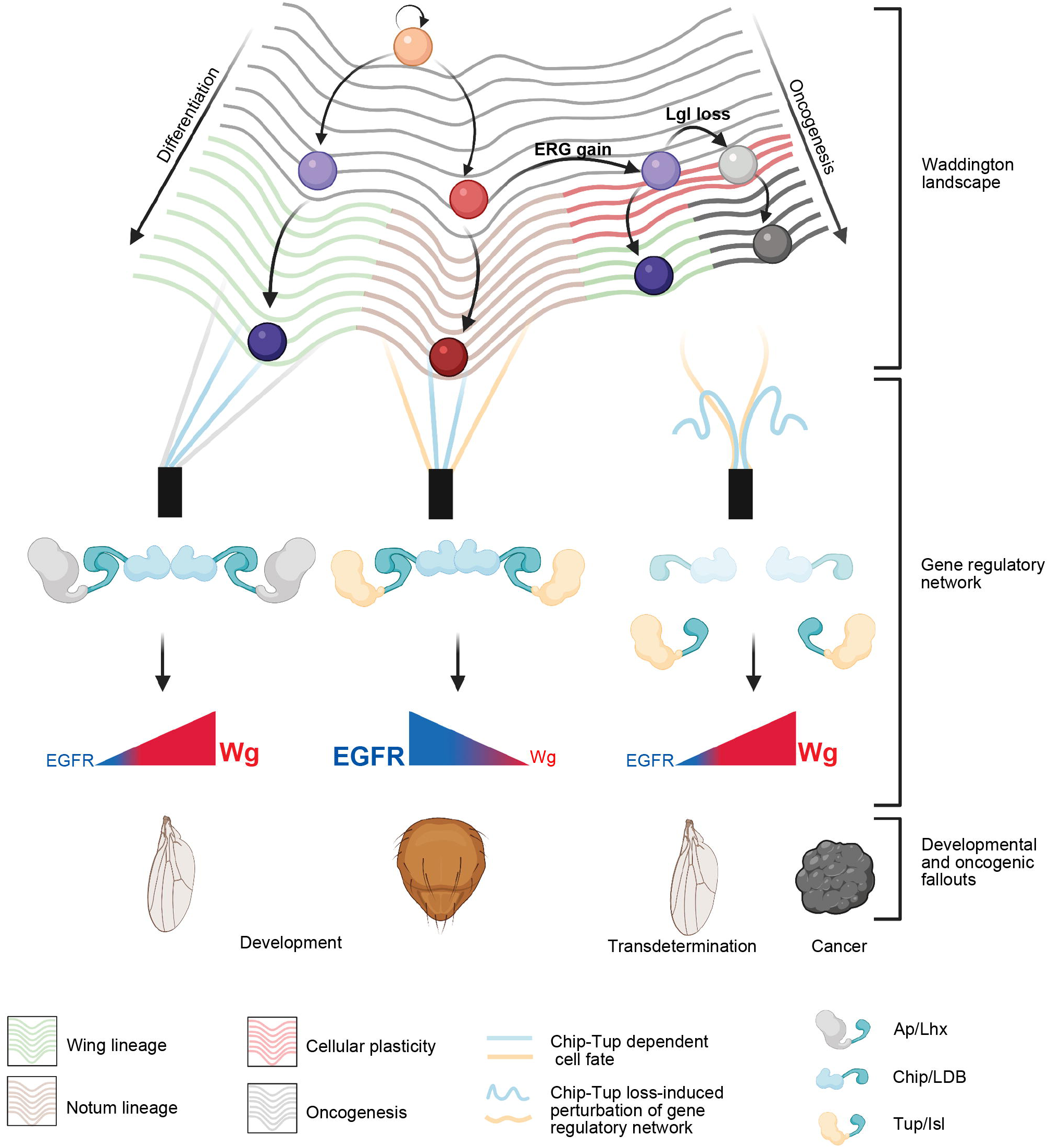
ERG-induced oncogenesis via induction of cellular plasticity in a Waddington field. A figurative Waddington landscape of a developing wing primordium. Pink ball at the top represents a cell with stem cell-like attributes. Pegs (represent individual signaling hubs) while guy-wires (represent the signaling outputs from these signaling hubs) that to valleys of notum versus wing cell fate, guided by the signaling outputs through the Chip-Tup (in this case), LIM-HD tetrameric complexes promoting either N-Wg (wing) or EGFR signaling (notum). ERG-mediated repression of Chip in the notum reactivates its N-Wg signaling. Two simultaneous oncogenic lesions (ERG gain and Lgl loss) tee the balls off from the valley of plasticity to that of oncogenesis.

In Waddington’s landscape, spatiotemporal recruitment of transcription factors creates a unique cell fate code (Moris et al., 2016). Oncogenic hits like ERG gain may induce stochastic fluctuations, meaning transcriptional noise, including non-mutational epigenetic reprogramming (Hanahan, 2022) that may culminate in loss of homeostatic balance of chromatin (Flavahan et al., 2017). These changes alter (Eldar and Elowitz, 2010; Raj and van Oudenaarden, 2008) Waddington’s landscape geometry (Sisan et al., 2012), marked by reversal to developmental ground states in oncogenically targeted cells (Khan et al., 2013). Our findings thus restate the maxim that carcinogenesis is cellular identity (Berdasco and Esteller, 2010) or development (Soto et al., 2008) gone awry.

### Future outlook

An *in vivo* approach to discover oncoprotein targets based on phenotypic fallout represents their functional identification. This approach in *Drosophila* can thus fast forward revelation of underlying cellular perturbations and forecast the gene regulatory network in cancers. A particular advantage of this approach is a rational extension of the observation from *Drosophila* to humans, based on deep homology. Amenability of *Drosophila* to sophisticated chemical genetic approaches (Dar et al., 2012) in combination with the discovery of oncoprotein targets would thus fast forward cancer drug screen complementing those of pharmaceutical approaches.

## STAR*METHODS

Detailed methods are provided in the online version of this paper and include the following:

- KEY RESOURCES TABLE
- LEAD CONTACT AND MATERIALS AVAILABILITY
- EXPERIMENTAL MODEL AND SUBJECT DETAILS

◦ Fly Strain and Generation of Clones
- METHOD DETAILS

◦ Generation of Transgenic Flies
◦ Immunohistochemistry and imaging
◦ Real-time qRT-PCR analysis
◦ Chromatin Immuno Precipitation-qPCR analysis
◦ Chromatin Immuno Precipitation sequencing (ChIP-Seq) analysis
◦ TF binding analysis
◦ Gene expression analysis
- QUANTIFICATION AND STATISTICAL ANALYSIS
- DATA AND CODE AVAILABILITY

Supplemental information, SI (Star methods, materials and methods, and Fig. S1 to Fig.S6).

## Author Contributions

Conceptualization: BA, PS

Methodology: MB, AB, UR, NM

Investigation: MB, AB, BA, PS

Visualization: MB, AB

Funding acquisition: PS, AB, BA

Project administration: PS

Supervision: PS, BA, AB

Writing – original draft: MB, PS

Writing – review & editing: MB, AB, BA, PS

## Supplementary figures

**Fig. S1.**
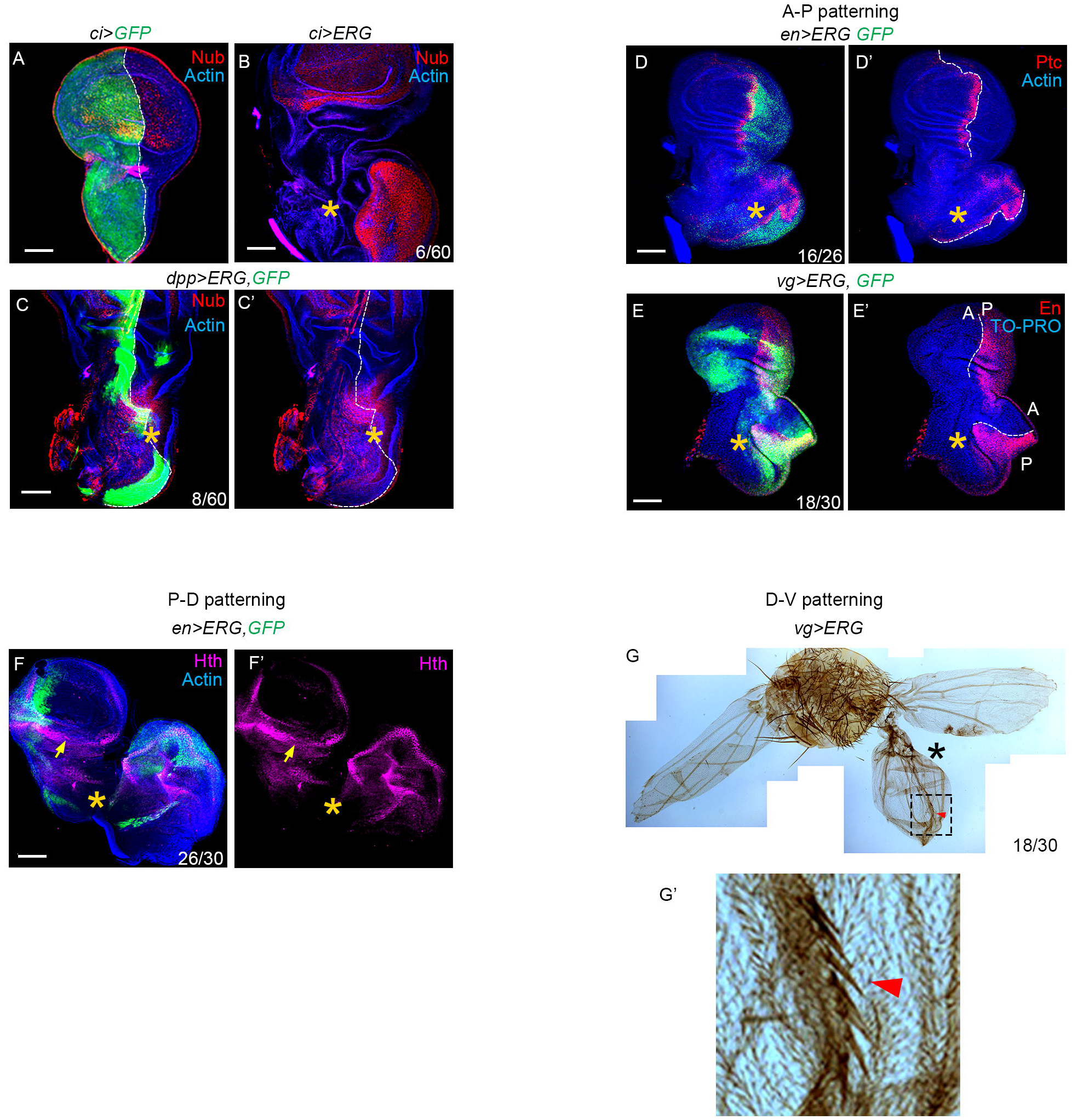
ERG-induced ectopic wing displays growth and patterning in all three axial coordinates. (**A-C**) Domain of *ci-Gal4* expression (*ci>GFP,* green, A). Broken line indicates it A-P boundary. Induction of Nub upon ERG expression in the notum (star, red, *ci>ERG*, B), apparently straddling past its presumed A-P boundary (see broken line in A). Expression of ERG under *dpp-Gal4* driver induces ectopic Nub straddling the A-P boundary (star, red C and C’, *dpp>ERG*). (**D, E**) *en>ERG* (D) and *vg>ERG* (E) wing imaginal discs. Broken line traces the A-P boundary of both endogenous and ectopic wing primordium (stars), lined by Ptc (red, D, D’) and En (red) and non-En compartments (E, E’), revealing their anterior-posterior (A-P) patterning. (**F**) *en>ERG* wing imaginal disc. Note the proximal Hth-expression in both endogenous (arrows) and ectopic wings (star), revealing their proximal-distal (P-D) patterning. (**G**) *vg>ERG* adult thorax mount displaying unilateral ectopic adult wing development (star, G). Boxed area in (G) is shown at higher magnification in (G’) to reveal triple row bristles along its D/V boundary, revealing it’s dorsal-ventral (D-V) patterning Scale bar-50µm; N= number of discs with desired marker or phenotype/ total wing discs observed

**Fig S2.**
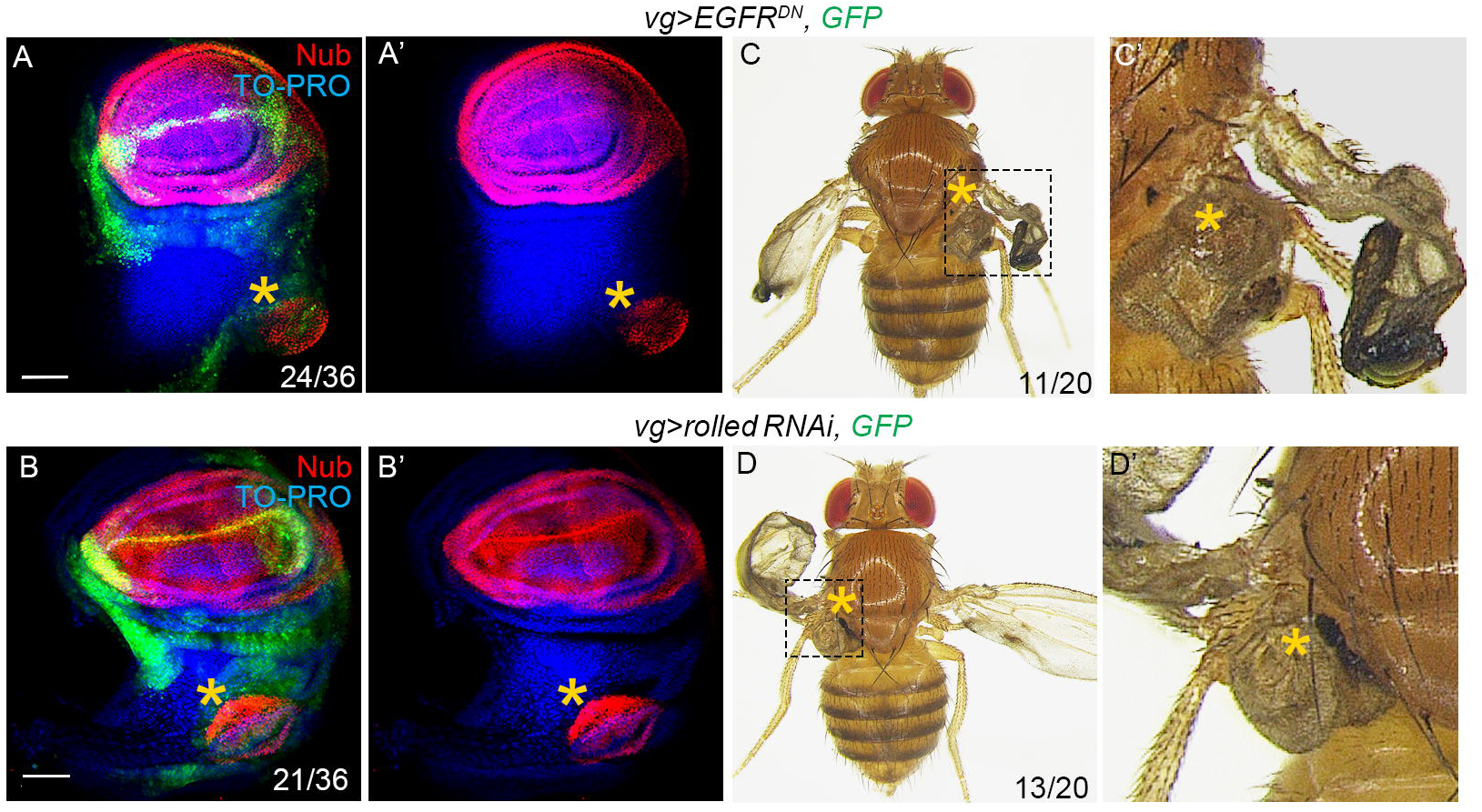
Loss of EGFR phenocopies ERG gain. (**A-D**) *vg>EGFR^DN^* (A, A’) or *vg> rolled-RNAi* (B, B’) display ectopic Nub in the notum, while their respective adults display development of ectopic wing from the mesothorax (C, C’-D, D’). Scale bar-50µm

**Fig. S3.**
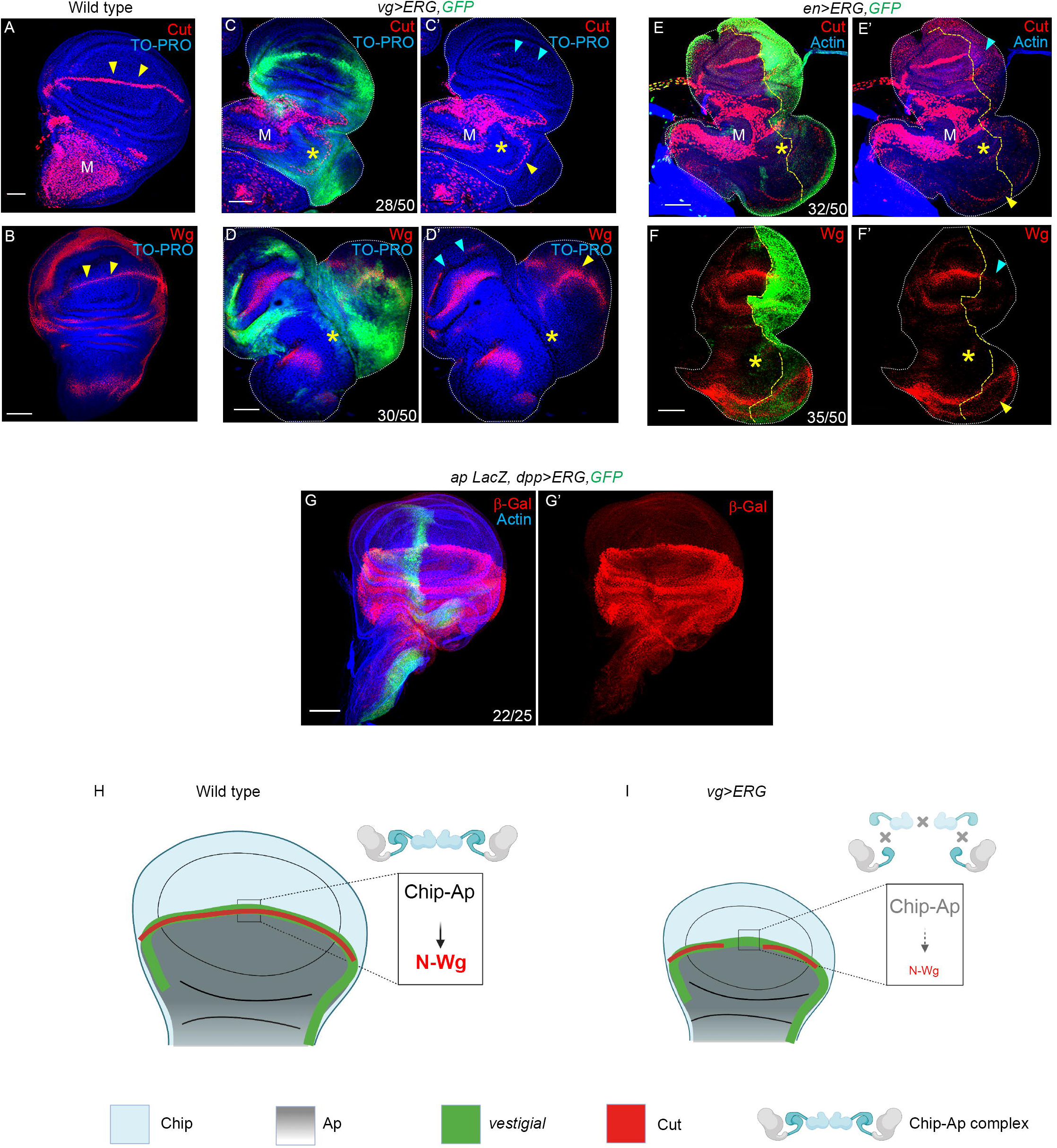
ERG gain suppresses growth and patterning of the endogenous wing. (**A, B**) Wild type wing imaginal discs displaying endogenous expression of N signaling target, Cut (red, A) and Wg (red, B) on the D/V boundary (yellow arrowhead). Note Cut expression in the notum (M) marks the adult muscle precursors (AMPs). (**C-F**) Expression of ERG under the *vg-Gal* driver (green, *vg>ERG*, *GFP*, C, D) and *en-Gal* driver (green, *vg>ERG*, *GFP*, E, F). Note the loss of Cut (blue arrowhead, C’, E’) and Wg (blue arrowhead, D’, F’) from the D/V margin of endogenous wing pouch in *vg>ERG* (C, D) and *en>ERG* (E, F) while these are seen in the D/V margins of their respective ectopic wing (yellow arrowhead, C’-F’). (**G**) *dpp>ERG* wing imaginal displaying unperturbed *ap-lacZ* reporter (anti-β-gal, red, G, G’). (**H, I**) Schematic representation of observations from (A-F). Chip-Ap mediated expression of N-Wg at D/V boundary (H) is compromised by gain of ERG (I). Scale bar-50µm

**Fig. S4.**
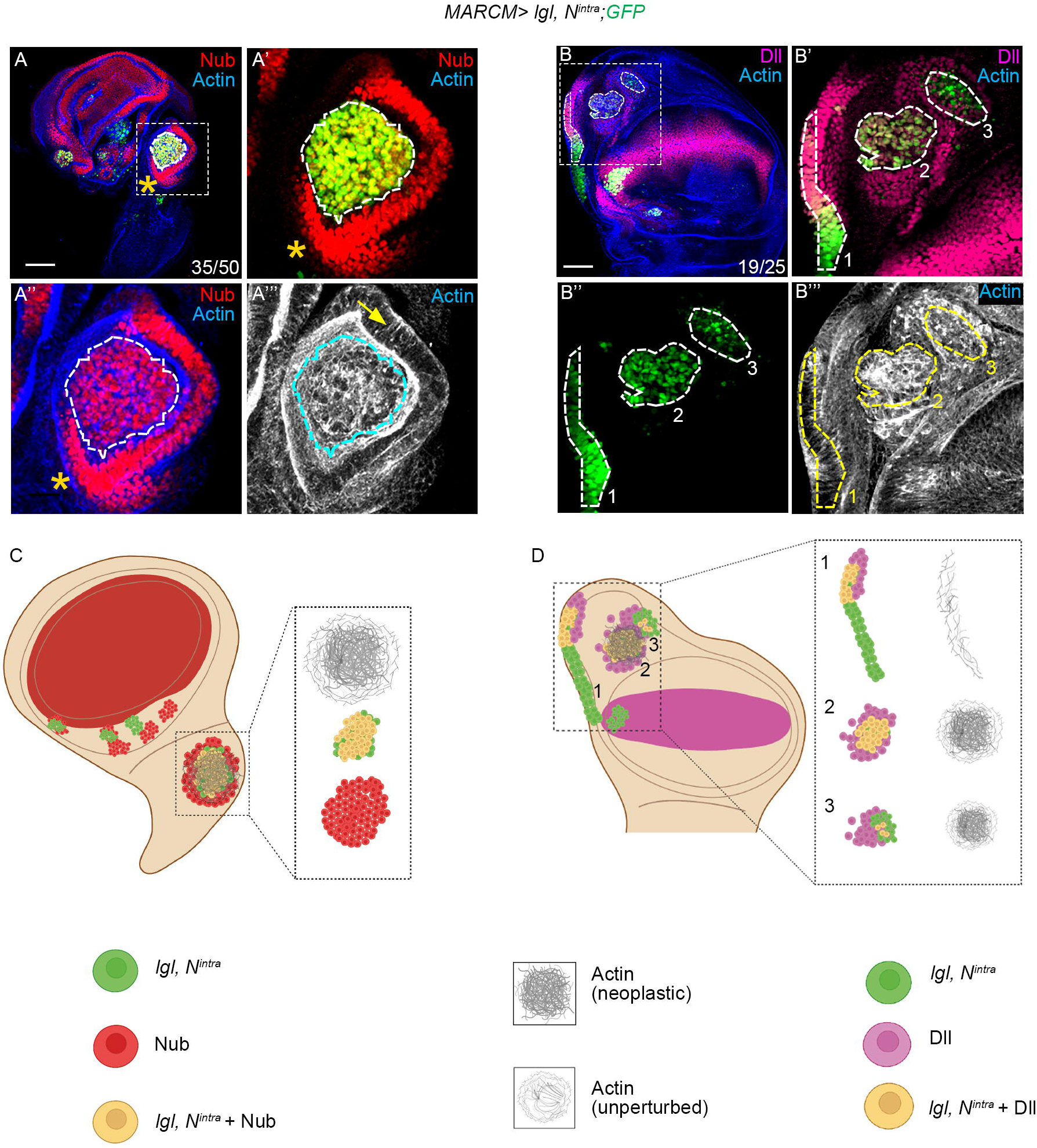
N-induced Wg morphogen drives lgl tumorigenesis in notum and hinge. (**A**) Notch-expressing *lgl* clones (*lgl; N^intra^*, GFP, green, A) in the notum display both autonomous (within the broken line) and non-cell-autonomous (outside broken line) induction of Wg target, Nub (star, red, A’- A’’). Boxed area in (A) is shown at a higher magnification in (A’, A’’, A”’). Note cell-autonomous neoplastic transformation of the clone (disrupted actin, grey) and hyperplasia in the surrounding (arrow, actin, gray, A”’). (**B**) *lgl; N^intra^* clones (GFP, green, B) in the inner fold of hinge. Boxed area in (B) is shown at a higher magnification in the rest of the panels (B’-B”’). Note the autonomous (within the broken line) and non-cell-autonomous (outside the broken line) induction of a Wg target, Dll in the clones marked as 1, 2 and 3 (B’-B’’’). Cell autonomous neoplastic transformation is seen only in the clone number 2 and 3, while clone 1 displays intact actin cytoskeleton (B’”). (**C, D**) Cartoon interpretations (C, D) of the results shown in (A, B), respectively. N-induced cell autonomous (C) and non-cell autonomous (D) activations of secreted Wg ligand targets, Nub (red) and Dll (magenta) in *lgl; N^intra^* clones in the notum and hinge domains of the wing imaginal discs. Scale bar- 50µm, N = number of clones displaying the characteristic phenotype out of the total scores in the notum (A) or hinge (B) domains.

**Fig. S5.**
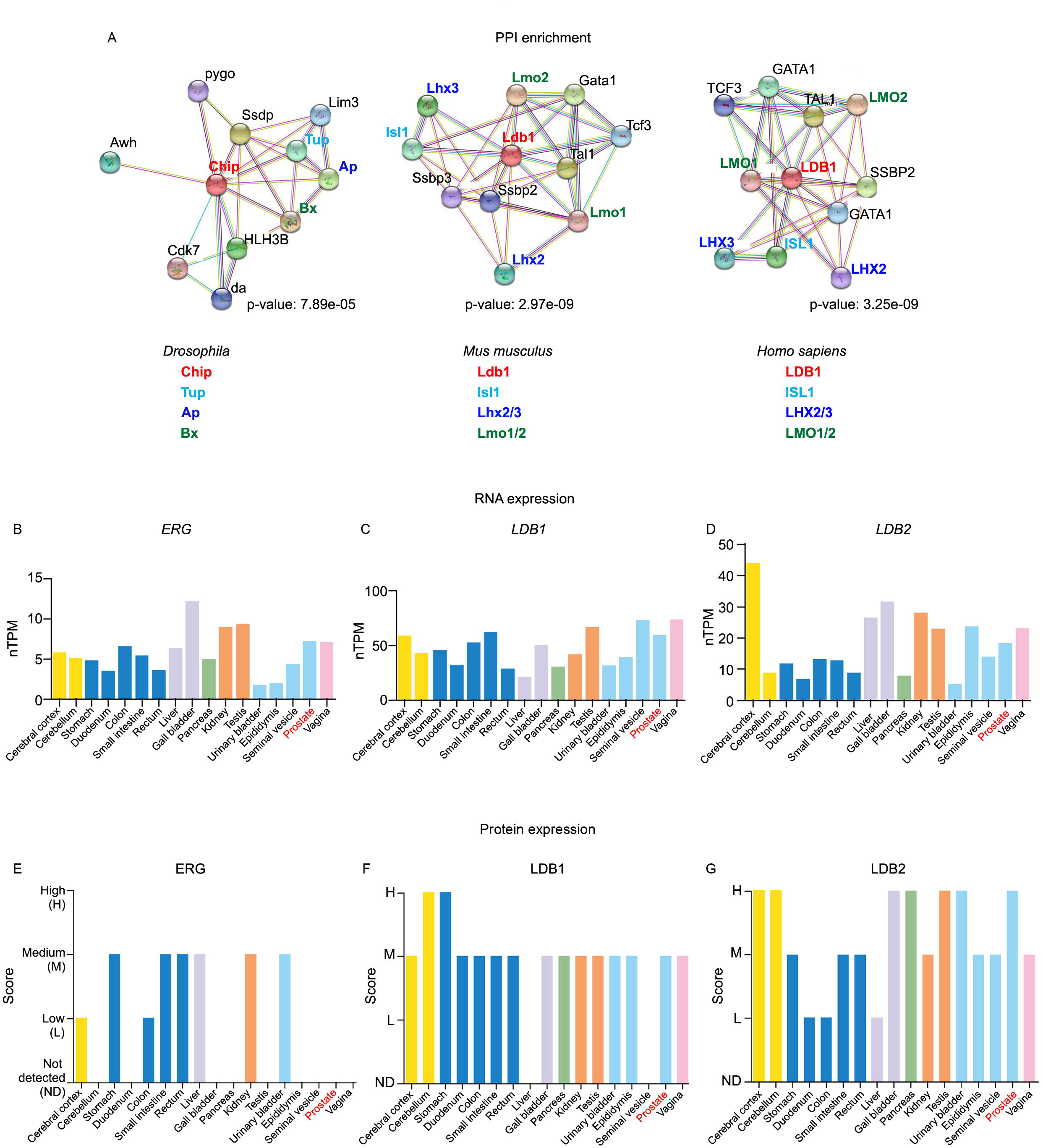
Conserved protein-protein interaction of Chip-Tup network and expression profile of ERG and LDB1 in healthy human tissues. (**A**) Protein-protein interaction (PPI) network using STRING (Szklarczyk et al., 2019) database of known (experimental) and predicted protein-protein interaction of direct (physical) and indirect (functional) associations. We noted a significant (P<0.001) and conserved interaction of Chip-Tup and LBD1-Isl1 protein in *Drosophila*, mice and human. (**B-G**) RNA (B-D) and protein (E-G) levels of ERG and LDB1 retrieved from The Human Protein Atlas (http://www.proteinatlas.org). RNA expression of *ERG* (B) is low, while *LDB1* (C) and *LDB2* (**D**) are moderate in the prostrate. By contrast, ERG is absent in prostate (E), while LDB1 (F) and LDB2 (G) protein expressions are medium and high, respectively. nTPM display number of transcripts per million for RNA expression and protein expression is graded from low (L), medium (M) to high (H).

---

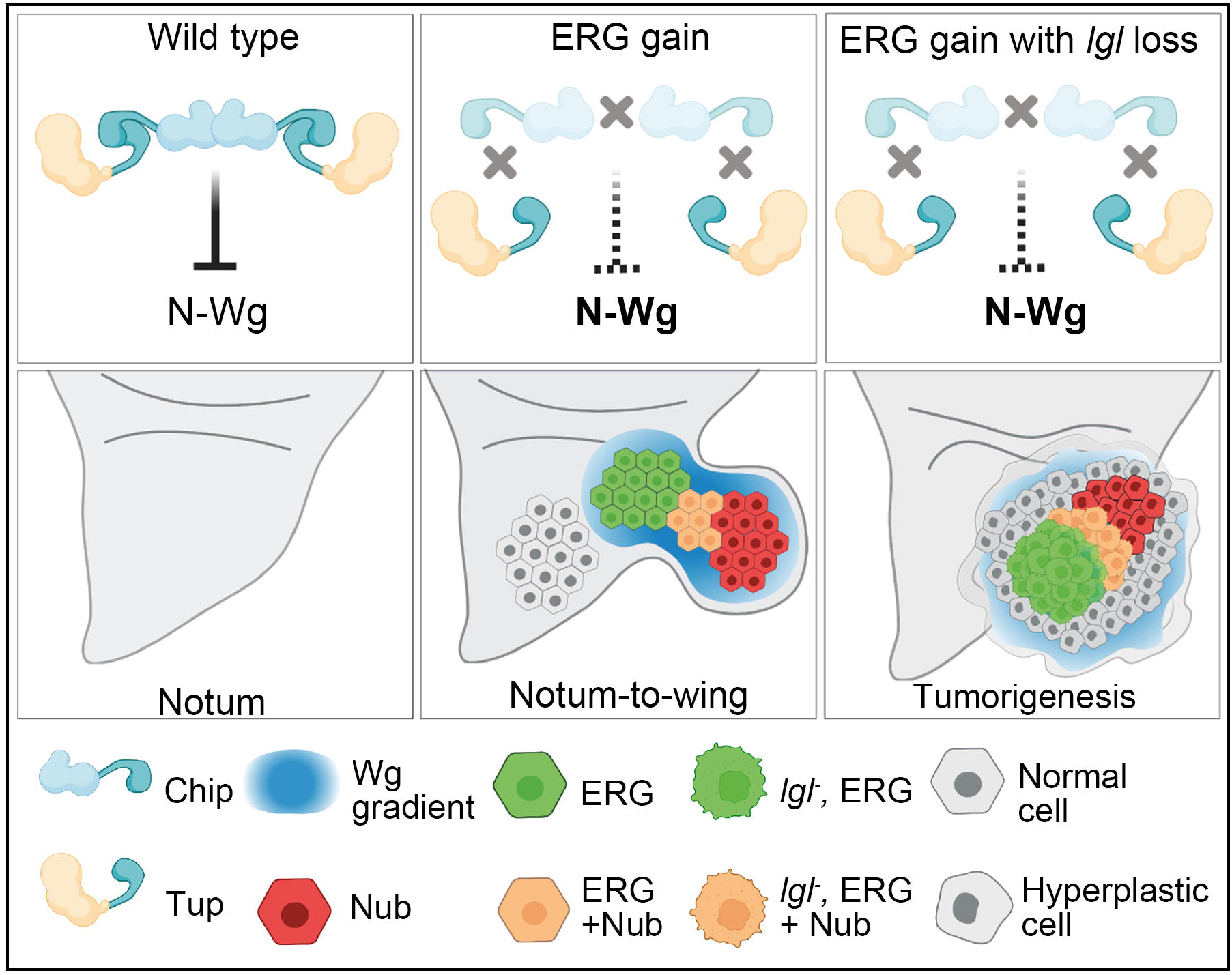

