## Supplementary Material and Methods for "Human ERG Oncoprotein Represses *Chip/LDB1* LIM-Domain Binding Gene in *Drosophila*"

**Supplemental information**

**STAR*METHODS**

**KEY RESOURCES TABLE**

| REAGENT OR RESOURCE | SOURCE | IDENTIFIER |
| --- | --- | --- |

**Antibodies**

Mouse Anti-Cut	DSHB	2B10

Mouse Anti-Nub	Gift from Steve Cohen,	------

Copenhagen, Denmark

Mouse Anti-Wg	DSHB	4D4

Mouse Anti-En	DSHB	4D9

Mouse Anti-Ptc	DSHB	Apa-1

Rabbit β-Gal	Sigma-Aldrich	SAB4200805

Rabbit Anti-PH3	Sigma-Aldrich	H0412

Mouse Anti-MMP1	DSHB	3B8, 5H7 and 23G1 (1:1:1)

Rabbit Anti-GFP	Invitrogen	A6455

Rabbit Anti-ERG	Abcam	ab92513

Rabbit Anti-EZH2	Abcam	ab191250

Rabbit anti-H3K27me3	Diagenode	pAb-069-050

Rabbit anti-IgG	Diagenode	RIG001

Goat Anti-Hth	Santa Cruz Biotechnology

Alexa Fluor 488 Goat anti-mouse IgG	Invitrogen	A32723

Alexa Fluor 555 Goat anti-mouse IgG	Invitrogen	A32727

Alexa Fluor 488 Goat anti-rabbit IgG	Invitrogen	A32731

Alexa Fluor 555 Goat anti-rabbit IgG	Invitrogen	A32732

Alexa Fluor 633 Goat anti-goat IgG	Invitrogen	A21086

TO-PRO-3	Invitrogen	S33025

Phalloidin-633	Invitrogen	A22284

ProLong™ Gold	Thermofisher	P36930

Experimental models: Organisms/Strain

*See Table S1*	N/A	N/A

Oligonucleotides

*See Table S2*	N/A	N/A

Software and Algorithms

Leica LAS AF software	Leica microsystem	<http://www.leicamicrosystems.com/productes/microscope-software/>

Prism	Graph pad	<https://www.graphpad.com/>

Excel	Microsoft	<https://www.microsoft.com/>

Photoshop	Adobe CS6	<http://www.adobe.com>

Illustrator	Adobe CC	<http://www.adobe.com>

BioRender	-	<https://biorender.com/>

JASPAR	-	<http://www.jaspar.genereg.net>

IGB	Integrated genome	<https://www.bioviz.org/>

browser

Other

Leica TCS SP5 microscope	Leica microsystem	[**http://www.leica-microsystems.com/**](http://www.leica-microsystems.com/)

**LEAD CONTACT AND MATERIALS AVAILABILITY**

Further information and requests for resources and reagents should be directed to and will be fulfilled by the Lead Contact, Pradip Sinha. In addition, *Drosophila* lines and other reagents generated in this study will be available upon request.

**EXPERIMENTAL MODEL AND SUBJECT DETAILS**

**Fly Strain and Generation of Clones**

*Drosophila* lines were cultured on standard yeast corn-meal agar food at 25^o^C. Egg-laying was carried out at 25^o^C for 4-6 hrs and, after 24 hours, shifted to 29^o^C to increase the efficiency of Gal4 activity (Duffy, 2002). Genetic screens were performed using the following tester strains: *vg Gal4* (Kim et al., 1996)*, en-Gal4* (Simmonds et al., 1995), *dpp-Gal4* (Lecuit et al., 1996), and *ci-Gal4* (gift from Ishwar Hariharan, UC Berkeley).

GFP-labelled mitotic clones were generated using flippase under heat shock promoter in imaginal discs of larvae (of the desired genotype). The detailed genotypes are provided at the end of the methods. flp/FRT-flip-out (Xu and Rubin, 1993) or MARCM (Lee and Luo, 2001, Khan et al., 2013) techniques were used; the latter also allowed simultaneous gain of a transgene by the UAS-Gal4 system. The eggs were laid in synchrony for 4-6 hrs; heat shock was given at 37°C for 30 minutes after 48 hrs of egg-laying. Imaginal discs were dissected from third instar larvae on day 3 after heat-shock (for control clones) and day 4 or 5 (for tumor clones).

**METHOD DETAILS**

**Generation of transgenic flies**

The transgenic flies expressing human *ERG* under UAS control were generated for this study. Briefly, cDNA encoding *ERG* (1.65kb was recloned from pCMV6 (6.6kb) vector (OriGene, Maryland, US) into P-element based pUAST (8.9kb) vector (Addgene, Cambridge, USA). The cDNA was amplified using PCR, purified MinElute kit, Qiagen, Hilden, Germany) was digested using BglII and NotI. The purified digested product (QIAquick gel extraction kit, Qiagen, Hilden, Germany). was ligated (T4 DNA ligase, NEB, US) into pUAST vector. Ultracompetent *E. coli* were transformed (One shot TOP10, *E. coli*, (Invitrogen, USA), using the ligation mix, subjected to selection (100µg/ml ampicillin). The positive clones were screened and confirmed via sequencing. Purified plasmid (QIAprep Spin Miniprep Kit, Qiagen, Hilden, Germany) were used to generate transgenic flies (Transgenic fly facility, C-CAMP, Bengaluru).

**Immunohistochemistry and imaging**

Larvae discs were dissected in cold 1xPBS and fixed in 1X PBS containing 4% formaldehyde and 0.2% Triton for 45 mins. Post 3 washes in PBS containing 0.2% Triton X-100, discs were incubated in primary antibody overnight at 4°C followed by blocking in 10% bovine serum albumin (90604-29-8), for 2 hours at room temperature, and subsequently with relevant fluorescent-tagged secondary antibody (1:250, Alexa fluorophore 488 or 555 or 633, Invitrogen). Discs were counter-stained with TO-PRO-3 (1:500) for nuclear stain or Phalloidin-633 (1:100) actin marker and mounted in anti-fade mounting media (ProLong™ Gold, Thermofisher #P36930). Samples were imaged with a 20x or 40x oil immersion objective on a Leica SP5 confocal microscope. Images were analyzed and processed using Leica confocal software-LAS AF and Adobe Photoshop CS-6.

**Real-time qRT-PCR analysis**

Total RNA was extracted from the VCaP PCa cells using the TRIZOL RNA extraction method (Sigma, T9424). 1.2 µg of total RNA from each sample was used to generate cDNA (SuperScript II Reverse Transcriptase, Thermo-scientific) using random primers (Macrogen). The resulting cDNA was used as a template for real-time qPCR with intron-spanning primers for *LDB1* and *GAPDH* (see Table S1 for primer sequence) on BIO-RAD using the SYBR Green qPCR master mix (Invitrogen). All assays were done in triplicate. RNA expression levels were normalized to GAPDH expression. *T*-tests were run on Graph Pad Prism 4.

**Chromatin Immuno-Precipitation-qPCR analysis**

Chromatin Immuno-precipitation (ChIP) was performed using LowCell# ChIP kit protein A (Diagenode, C01010072), following instructions to users. Briefly, approximately 300 wing discs were used per reaction, and these were dissected in ice-cold 1× PBS and crosslinked in 1% freshly prepared formaldehyde (Sigma-Aldrich #F15587) for 15 min at 37°C. Crosslinking was quenched with 1.25 mM freshly prepared glycine (LowCell# ChIP kit) for 5 min. The cells were lysed in 130µl Buffer B (LowCell #ChIP kit) supplemented with complete protease inhibitor (Roche #P5147) and phenylmethylsulfonyl fluoride (PMSF; Sigma-Aldrich, #P7626). 130 µl of the lysate was sheared using a Bioruptor (Diagenode) at high frequency for 15 cycles of 30 seconds ON and 30 seconds OFF. 870 µl of Buffer A (LowCell# ChIP kit) supplemented with complete protease inhibitor (Roche) and PMSF (Sigma-Aldrich) was added to the sheared chromatin; and 10 µl of which lysate was saved as input control, while the remaining was incubated with magnetic beads (11 µl, LowCell #ChIP kit) which were pre-incubated (4 hrs at 4oC) with the required antibodies: rabbit anti-ERG (Abcam, ab92513), or rabbit anti-GFP (Invitrogen, A6455), or rabbit anti-H3K27me3 (diagenode, pAb-069-050), or rabbit anti-EZH2 (Abcam, ab191250) or rabbit anti-IgG (diagenode, RIG001) overnight at 4^o^C. The beads-antibody complex was separated using a magnetic rack. The immobilized chromatin was then eluted in 100 µl DNA elution buffer (1% SDS, 0.1 M sodium bicarbonate with proteinase K and RNaseA). ChIP eluates and input (10%) were assayed for the enrichment of the desired DNA fragment using real-time PCR using SYBR Master Mix (Invitrogen) and oligomers to desired genes (**table S2**).

**Chromatin Immuno-Precipitation Sequencing (ChIP-Seq) analysis**

To determine the recruitment of ERG on *LDB1* and *LDB2* promoter, we analyzed publicly available ChIP-Seq data (GSE28950) for ERG and treatment with siERG (GSE116055) in VCaP cells. ChIP-Seq peaks for ERG and siERG treated samples were called using Model-based analysis of ChIP-seq (MACS; *P* < 10^−5^) with default settings against the input. BAM and BED files obtained were visualized by Integrative Genomic Browser (IGB).

**TF binding analysis**

The presence of putative ERG binding sites (EBS) in *E(z)* and *Chip* promoters, and E(z) binding sites, namely the Polycomb Response Elements (PREs), in the *Chip* promoter region were scanned using publicly available transcription factor binding prediction software, JASPAR.

**Gene expression analysis**

Differentially regulated genes were analyzed in publicly available datasets of ERG-overexpressing benign RWPE (GSE86232) and VCaP cells upon ERG knockdown (GSE110656 and GSE35540) retrieved from Gene Expression Omnibus (GEO) database with an accession number. In addition, in both the cell lines, the microarray data set were analyzed for *LDB1* expression.

**Quantification and Statistical Analysis**

***Percent wing disc displaying ectopic wing***

Wing discs with ectopic Nubbin phenotype (%) were calculated by analyzing ectopic wing Nub expression/total number in at least 20 wing epithelia/set (total N=60) of each Gal4 (*en-Gal4, vg-Gal4, dpp-Gal4, and ci-Gal4*). Statistical analysis was performed using Excel and Graph pad prism, and data were analyzed by one-way ANOVA that plotted with mean ± SEM displaying a significance value *p*<0.0001.

***Percent flies displaying ectopic wing***

From a set of 100 adults each, wherein *ERG* (UAS-*ERG)* was driven with *vg-Gal4* (*vg> ERG*), the percent of eclosed adults with and without ectopic wings were calculated for three such replicates. Statistical significance was computed using an unpaired t-test and plotted mean ± SEM displaying a significance value *p*<0.01.

**Data and code availability**

Not applicable

**Detailed genotypes of fly strain used in the study (relevant genotypes are marked in boldface)**

*Figure 1*

***en>GFP:*** *w;* *en-Gal4*; *UAS GFP/+*

***nub>mCherry****: w; nub-Gal4, UAS mCherry*

***vg>ERG, GFP****: w;* *vg-Gal4*; *UAS GFP/ UAS ERG*

***en>ERG, GFP****: w;* *en-Gal4*; *UAS GFP/ UAS ERG*

***MARCM>ERG*** clone*: y w hs-flp tub Gal4 UAS-GFP; FRT40/tub-Gal80 FRT 40; UAS-ERG*

*Figure 2*

***vg>N^intra^, GFP:*** *w; vg-Gal4/ UAS N^intra^; UAS GFP/+*

***en>N^DN^; ERG, GFP:*** *w; en-Gal4/ UAS N^DN^; UAS GFP/UAS ERG*

***en>mam^DN^; ERG, GFP:*** *w; en-Gal4/ UAS mam^DN^; UAS GFP/UAS ERG*

***en>GPI dFz2; ERG, GFP:*** *w; en-Gal4/+; UAS GFP/UAS ERG, UAS GPI dFz2*

*Figure 3*

***hs flp act> Chip^Δoid^*** clones***:*** *w hs-flp; act>y+> Gal4 UAS GFP/+; UAS* *Chip^∆oid^ /+*

***en>Chip^Δoid^, GFP :*** *w; en-Gal4/+; UAS Chip^Δoid^/UAS GFP*

***vg>Chip^Δoid^ :*** *w; vg-Gal4/+; UAS Chip^Δoid^/+*

***en>Chip; ERG, GFP:*** *w; en-Gal4/UAS Chip; UAS ERG/UAS GFP*

***vg>Chip; ERG:*** *w; vg-Gal4/UAS Chip; UAS ERG/+*

***en>tup RNAi, GFP*:** *w; en-Gal4/+; UAS tup RNAi /UAS GFP*

***vg>tup RNAi*:** *w; vg-Gal4/+; UAS tup RNAi /+*

***en> tup; ERG, GFP:*** *w; en-Gal4/UAS Tup; UAS ERG/UAS GFP*

*Figure 4*

***E(z)-GFP, en>ERG:*** *w; en-Gal4-UAS RFP/E(z)-GFP; UAS ERG/+*

*Figure 5*

Genotypes of MARCM clones

***lgl^-^; ERG:*** *y w hs-flp tub Gal4 UAS GFP; lgl^4^ FRT40/tub-Gal80 FRT 40; UAS ERG/MKRS*

***lgl^-^, Chip; ERG:*** *y w hs-flp tub Gal4 UAS GFP; lgl^4^ FRT40 UAS Chip/tub-Gal80 FRT 40; UAS ERG/MKRS*

***lgl^-^, E(z) RNAi; ERG:*** *y w hs-flp tub Gal4 UAS GFP; lgl^4^ FRT40 UAS Ez RNAi/tub-Gal80 FRT 40; UAS ERG/MKRS*

***lgl^-^; GPI dFz2, ERG:*** *y w hs-flp tub Gal4 UAS GFP; lgl^4^ FRT40 /tub-Gal80 FRT 40; UAS GPI Dfz2, UAS ERG/MKRS*

Figure S1

***ci>GFP****: w; ci-Gal4/+; UAS GFP/+*

***ci>ERG****: w; ci-Gal4/+; UAS ERG/+*

***vg>ERG, GFP****: w; vg-Gal4/+; UAS GFP/UAS ERG*

***dpp>ERG, GFP****: w;*; *dpp-Gal4*-*UAS GFP/ UAS ERG*

Figure S2

***vg>EGFR^DN^, GFP:*** *w; vg-Gal4/UAS EGFR^DN^; UAS GFP/+*

***vg>rolled RNAi, GFP:*** *w; vg-Gal4/UAS rolled RNAi; UAS GFP/+*

Figure S3

***ap LacZ, dpp> ERG, GFP:*** *w;; dpp-Gal4, UAS GFP/ap LacZ, UAS ERG*

Figure S4

***lgl^-^; N:*** *y w hs-flp tub Gal4 UAS GFP; lgl^4^ FRT40/tub-Gal80 FRT 40; UAS N*

**Table S1: Source of fly lines**

*UAS-ERG*	This study

*UAS-N^intra^*	BDSC

*UAS-N^DN^*	BDSC

*UAS-mam^DN^*	BDSC

*UAS-GPI-dFz2*	BDSC

*UAS-Chip^Δoid^*	BDSC

*UAS-Chip*	BDSC

*UAS-tup*	BDSC

*UAS-tup RNAi*	BDSC

*ap-LacZ*	BDSC

*UAS-E(z) RNAi*	BDSC

*UAS-EGFR^DN^*	BDSC

*UAS-rl RNAi*	BDSC

*lgl FRT40*	Khan et al, 2013

**Table S2: List of primers used in RT-PCR and ChIP-PCR**

| **q-PCR sequences** | | **Forward primer (5'🡪3')** | **Reverse primer (5'🡪3')** |
| --- | --- | --- | --- |
| *GAPDH* | | TGCACCACCAACTGCTTAGC | GGCATGGACTGTGGTCATGAG |
| *LDB1* | | ACGACGAGGACAGCTTTAAC | GGTTCTCCGATTTGCTTTCTTG |
| **ChIP-PCR sequences** | | **Forward primer (5'🡪3')** | **Reverse primer (5'🡪3')** |
| ERG binding on *LDB1* promoter **(EBS)** | | ACACCACTGTGTCCAACTAAC | GGCCAGCCAGAAGTTCAATA |
| ERG binding on *E(z)* promoter **(EBS)** | | CAAACCCAGTTATGGTTGGTAAAC | CAGCCTGCGAAGGTATGTAA |
| ERG binding on *Chip* promoter | **EBSI** | CCGAGTTAGCCTTTCTTTCCT | GACGGAGGCGTGGTAAATAG |
|  | **EBSII** | AGCCCTTGGAAATAAGCAGAG | CATCTTCCAGTGTCCGTTGAA |
| E(z) binding on *Chip* promoter | **PRE-I** | AATCGATTAAATACGACAACT CCAG | GCAGCCAACACTCACTCATA |
|  | **PRE-II** | CGATGAGAAATCCAGCGACTAT | CGAGCTTTCTGAACCCAATCT |
|  | **PRE-III** | TTCCTGTGCAGTTCCTTGTC | GTGTGCGTTTCCTACACTCTAA |
